# Natural and patient-derived mutations in BK polyomavirus VP1 reveal structural determinants of BC-loop dependent antibody escape

**DOI:** 10.1101/2025.08.09.669497

**Authors:** Onno Akkermans, Cristina Ubeda Nicolau, Asanga Bandara, Consuelo C. Sierra, Fernando H. Martins, Steff De Graef, Erin E. Deans, Stephen J. Ross, Senne Sienaert, Miriam Cerrada, Vipul C. Chitalia, Nagraj Mani, Stephen D. Weeks, Ali H. Munawar

## Abstract

BK polyomavirus (BKPyV) reactivation poses a serious threat to both renal and hematopoietic transplant outcomes, with no approved antiviral therapies. Neutralizing monoclonal antibodies targeting the viral capsid protein VP1 have recently advanced into clinical development, driven by promising *in vitro* potency. Since naturally elicited neutralizing antibodies arise following infection, we examined whether host immune pressure may have shaped the VP1 surface, with implications for therapeutic application. To probe the impact of VP1 coevolution on therapeutic engagement, we evaluated two neutralizing antibody formats: the clinical-stage monoclonal antibody 319C07 and nanobodies VHH16 and VHH17. Our X-ray crystallographic analysis of the VP1 pentameric core in complex with antibody fragments showed a convergent receptor-mimetic engagement of the sialic acid-binding cleft, mapping to a shared epitope centered around the VP1 BC-loop. Analysis of over 900 BKPyV VP1 sequences from public databases, combined with longitudinal VP1 sequence data from transplant recipients, highlighted widespread pre-existing diversity at the BC-loop contact sites, underscoring the evolutionary adaptability of BKPyV at this antigenic surface that may compromise therapeutic recognition. Through integrated mutagenesis and binding analyses, we show that circulating BC-loop mutations, including single amino acid substitutions, are sufficient to abrogate 319C07 binding and neutralization. Nanobodies VHH16 and VHH17, selected for their compact size and potential to access intrarenal sites of BKPyV replication, also exhibited complete loss of binding across multiple clinically observed VP1 variants. Our findings may offer a mechanistic framework to interpret the translational gap between *in vitro* neutralization and clinical efficacy of these biologics. As part of ongoing efforts to target VP1, we demonstrate, for the first time, small molecules with nanomolar to sub-nanomolar affinities against BKPyV VP1 variants including potent binding to JCPyV VP1. Our work paves a path towards a new class of antiviral strategies that could block replication at intracellular replication sites, provide a higher genetic barrier to resistance and the potential to address both viral reactivation and persistent, high-burden viremia in transplant settings.

## Introduction

BK polyomavirus (BKPyV) is a near-ubiquitous virus that establishes a lifelong latent infection in the urothelium during early years of life [1]. Seroprevalence exceeds 80% in adults worldwide, with reactivation occurring in immunocompromised individuals, most notably following kidney transplantation [2, 3]. Up to 30-40% of kidney transplant recipients develop viruria, with a subset progressing to viremia and BKPyV-associated nephropathy (BKPyVAN), a destructive tubulointerstitial disease that leads to irreversible allograft dysfunction and loss [2, 4]. This clinical trajectory is especially consequential given the scarcity of donor organs, the protracted wait times for transplantation in end-stage renal disease patients, and the intensive surveillance, immunosuppressive management, and the risk of dialysis associated with graft failure [5, 6]. Beyond renal transplantation, BKPyV reactivation is frequently observed following hematopoietic stem cell transplants (HSCT), the second most common transplant procedure globally, where viruria affects up to ∼50% of recipients, leading to hemorrhagic cystitis in as many as ∼55% [7]. Reactivation is also increasingly recognized in lungs, heart, and liver recipients, with prevalence estimates ranging from 10% to 30%, depending on immunosuppressive regimen intensity and diagnostic assay sensitivity [8]. The broader impact of BKPyV across solid organ and hematopoietic transplants has historically been obscured by inconsistent diagnostics and variable clinical awareness. The recent FDA clearance of a standardized quantitative PCR assay (Roche Cobas® BK Virus Test, 2020) has marked a turning point in surveillance and clinical recognition. Yet despite this progress, no targeted antiviral therapies currently exist. In clinical practice, current management of BKPyV reactivation relies on modulation of the host immune response, often through empiric reduction of immunosuppression, a strategy that carries risk of precipitating allograft rejection [6].

The absence of effective antiviral therapies for BKPyV is largely attributable to the genomic constraints of the virus and lack of classical drug targets. BKPyV shares its genome organization with other human polyomaviruses, including JC polyomavirus (JCPyV), which infects glial cells [9] and causes progressive multifocal leukoencephalopathy (PML) in immunosuppressed patients receiving immunomodulatory biologics for autoimmune disease and transplantation [10, 11]. Unlike many pathogenic DNA and RNA viruses, polyomaviruses like BKPyV and JCPyV, lack virally encoded enzymes that have historically served as the molecular targets for classical small-molecule antiviral drug design. This shared architecture highlights a defining challenge of polyomavirus biology: the absence of classical viral drug targets and reliance on host cell machinery instead. Its ∼5 kilobase circular, double-stranded DNA genome encodes a compact set of early regulatory and late structural proteins, including the large and small tumor antigens (LTAg and sTAg), the agnoprotein, and three capsid proteins: VP1, VP2, and VP3 [12]. Among these, LTAg and VP1 are essential for the viral life cycle. Efforts to target the ATP-dependent helicase domain of LTAg, failed to yield viable compounds [13]. BKPyV VP1, the major capsid protein forms the icosahedral shell of the unenveloped virion through the assembly of 72 pentamers and furnishes the dominant antigenic surface [14]. Antibodies targeting BKPyV VP1 have seen clinical advancement in recent years (**Figure 1A and 1B**) [15, 16].

**Figure 1:**
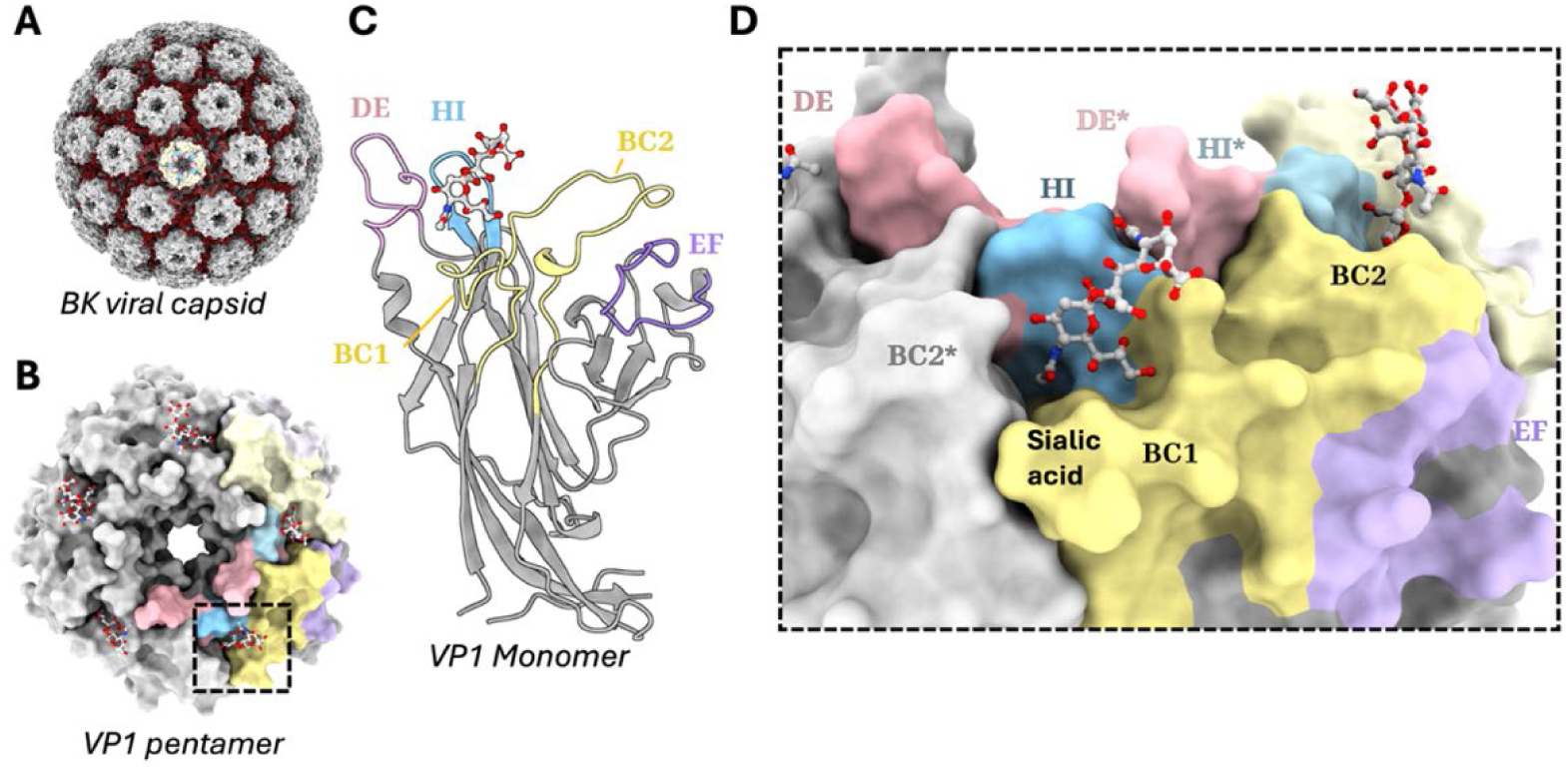
Surface exposed loops on BKPyV VP1 form the host receptor binding pocket of the viral capsid. **A)** Surface representation of BKPyV viral capsid (PDB: <u>6GG0</u>) **B)** Surface representation of the VP1 pentamer bound to sialic acid molecules (PDB: <u>4MJ0</u>). VP1 is colored grey, while the loops forming the sialic acid binding pocket of two monomers are colored yellow (BC position 55-85, pink (DE positions 132-142), purple (EF positions 167-181), blue (HI positions 270-277). and **C)** The VP1 monomer is depicted in cartoon representation with the loops colored (PDB: <u>4MJ0</u>). **D)** Close up of the sialic acid binding site. Asterisk represents loops that are from a neighboring monomer that contribute to the binding pocket.

These features position VP1 as both a structural linchpin of viral entry and a focal point of host immune recognition. Four serotypes (I-IV) of BKPyV are classified based on VP1 sequence divergence, with type I predominating globally and types II-IV exhibiting regional enrichment [12].

Serological surveys indicate that by early adulthood, most individuals harbor anti-BKPyV IgGs from childhood infection [17]. Majority of these antibodies are non-neutralizing, acting via Fc-mediated effector functions, although their contribution is not well defined clinically [18]. A critical fraction of the humoral response comprises neutralizing antibodies that engage surface-exposed epitopes on VP1 via their Fab domains, preventing capsid attachment and viral entry. Neutralizing antibodies act independently of T-cell-mediated immunity, making them particularly relevant for viral control in immunosuppressed transplant recipients. In transplant cohorts, pre-transplant neutralizing antibody titers below 4 log_10_ portend a nearly two-fold higher risk of viremia, whereas titers above 4 log_10_ halve that risk [19]. Moreover, passive transfer of BKPyV-neutralizing intravenous immunoglobulin (IVIG) in low titer patients has been shown to recapitulate the protection observed in naturally high-titer individuals [20]. These clinical observations underscore neutralizing antibodies as a correlate to BKPyV control and provide a compelling rationale for the development of VP1-targeted monoclonal antibodies.

Among emerging efforts to harness the therapeutic potential of neutralizing antibodies, 319C07 (Potravitug), a highly potent human IgG monoclonal antibody is currently under clinical evaluation for transplant-associated BKPyV infection [16]. Potravitug and other antibody-based approaches such as MAU868 [15] exploit the fact that VP1 is the sole surface-exposed structural protein of BKPyV and the primary target of the humoral immune response.

The VP1 surface comprises five solvent-exposed loops, BC, DE, EF, GH, and HI (**Figure 1C).** [21]. Among these, the BC-loop has emerged as the immunodominant region, recurrently targeted by antibodies elicited through natural infection. Seroprofiling of transplant recipients uncovers a strong bias toward BC-loop-reactive antibodies, and longitudinal viral sequencing reveal an elevated nonsynonymous-to-synonymous substitution ratio within BC-loop codons, consistent with antibody-driven selection *in vivo* [22]. Beyond its antigenicity, the BC-loop also forms part of the receptor-binding surface and contributes to sialylated glycan engagement, positioning it at the intersection of immune pressure and viral entry [23]. While the structural footprints of some therapeutic antibodies in complex with VP1 have been resolved, it remains unclear how naturally occurring sequence variation within the BC-loop modulates antibody binding and efficacy.

Inspired by these observations and the advent of clinical-stage antibodies targeting this region, we set out to define the structural and mechanistic basis of BC-loop recognition, and how naturally occurring variation modulates these interactions. We report X-ray structures of 319C07, and two camelid derived single-domain (heavy chain) antibodies, ORT-VHH16 (VHH16) and ORT-VHH17 (VHH17), bound to the BKPyV VP1 pentamer. These nanobodies were selected for their potent *in vitro* neutralization activity. Owing to their small size and compact paratope architecture, nanobodies can access sterically restricted epitopes that conventional IgGs cannot and are amenable to multivalent or bi-paratope formats. Crucially, their ability to traverse the glomerular filtration barrier makes them uniquely suited for targeting BKPyV in the renal tubular epithelium, the exclusive site of viral replication. Through comparative structural analysis, we establish distinct antibody binding modes across the VP1 surface. In parallel, we perform a comprehensive bioinformatic survey of BC-loop variation and functionally assess the impact of prevalent mutations using a combination of binding and pseudovirus neutralization assays. These data illuminate how the evolutionary and structural plasticity of the BC-loop enables immune escape and underscore the challenge of achieving broad and durable control through antibody-based strategies alone.

## Results

### A. Clinical-stage anti-BKPyV antibody 319C07 engages the receptor-binding cleft on VP1 through topological mimicry of Neu5Ac

To define the molecular mechanism of therapeutic neutralization of 319C07, we determined the X-ray crystal structure of the Potravitug single-chain variable fragment (scFv-319C07) bound to the BKPyV VP1 pentamer at 4.1 Å resolution (**Table 1**).

**Table 1:** Data collection and refinement statistics

| Protein | BK VP1 (Genotype 1a) | BK VP1 (Genotype 1a) | BK VP1 (Genotype 1a) |
| --- | --- | --- | --- |
| scFv / VHH | 319C07 | VHH16 | VHH17 |
| Beamline | ALBA | Proxima 2 | Proxima 2 |
| **Data Collection:** |  |  |  |
| Space group | P2<sub>1</sub> 2<sub>1</sub> 2<sub>1</sub> | P1 2<sub>1</sub> 1 | P1 2<sub>1</sub> 1 |
| **Cell dimensions:** |  |  |  |
| a, b, c (Å) | 110.3, 151.9, 164.7 | 117.6, 135.3, 167.8 | 85.8 152.4 91.9 |
| $\alpha$, $\beta$, $\gamma$ (°) | 90, 90, 90 | 90, 97.8, 90 | 90, 94.4, 90 |
| Resolution (Å) | 75.9-3.77 (3.79-3.77) | 116-2.59 (2.64 – 2.59) | 47.3- 2.49 (2.52-2.49) |
| R<sub>p.i.m</sub> (%) | 2.3 (123) | 3.5 (146) | 10.4 (92.5) |
| I/$\sigma$ | 3.7 (0.7) | 5.1 (0.5) | 6.5 (0.7) |
| CC1/2 | 1.00 (0.4) | 0.98 (0.33) | 0.99 (0.29) |
| Completeness (%) | 99.9 (100) | 99.9 (99.9) | 99.3 (80.1) |
| Redundancy | 8.5 (8.9) | 6.3 (5.9) | 3.6 (3.1) |
| **Refinement:** |  |  |  |
| No. reflections | 29566 (1460) | 160773 (5239) | 81437 (2274) |
| No. free reflections | 1473 (131) | 8023 (260) | 4072 (113) |
| R<sub>work</sub>/R<sub>free</sub> | 0.257 / 0.309 (0.398 / 0.386) | 0.203 / 0.245 (0.337 / 0.382) | 0.191 / 0.257 (0.359 / 0.394) |
| **No. atoms:** |  |  |  |
| Protein | 15318 | 29406 | 14951 |
| Water | 0 | 195 | 969 |
| **B factors (Å<sup>2</sup>):** |  |  |  |
| Protein | 43.55 | 55.14 | 45.48 |
| Water | N/A | 42.49 | 50.50 |
| **RMSDs:** |  |  |  |
| Bond lengths (Å) | 0.009 | 0.009 | 0.009 |
| Bond angles (°) | 1.77 | 1.11 | 1.53 |
| **Ramachandran:** |  |  |  |
| Favored (%) | 91.5 | 96.17 | 94.02 |
| Outlier (%) | 1.24 | 0.16 | 0.89 |
| Note: Values in parentheses reflect the highest resolution shell. |  |  |  |

The antibody engages the apical face of the VP1 pentamer, bridging two adjacent monomers (**Figure 2A** **and** **Figure 3A**). Its VH domain penetrates deeply into the receptor-binding cleft, while the VL domain extends laterally across the BC-loop creating a bi-partite interaction surface. 319C07 anchors over the BC-loop hairpin of one monomer (BC2) and contacts adjacent receptor-binding surfaces, including BC1 (e.g., K69) and the HI-loop (residues N270-T277) of a neighboring monomer. The electron density map unambiguously resolved the antibody binding footprint and orientation relative to the sialic acid receptor binding site (**Figure 3B and 3C**) providing a robust basis for structural interpretation.

**Figure 2:**
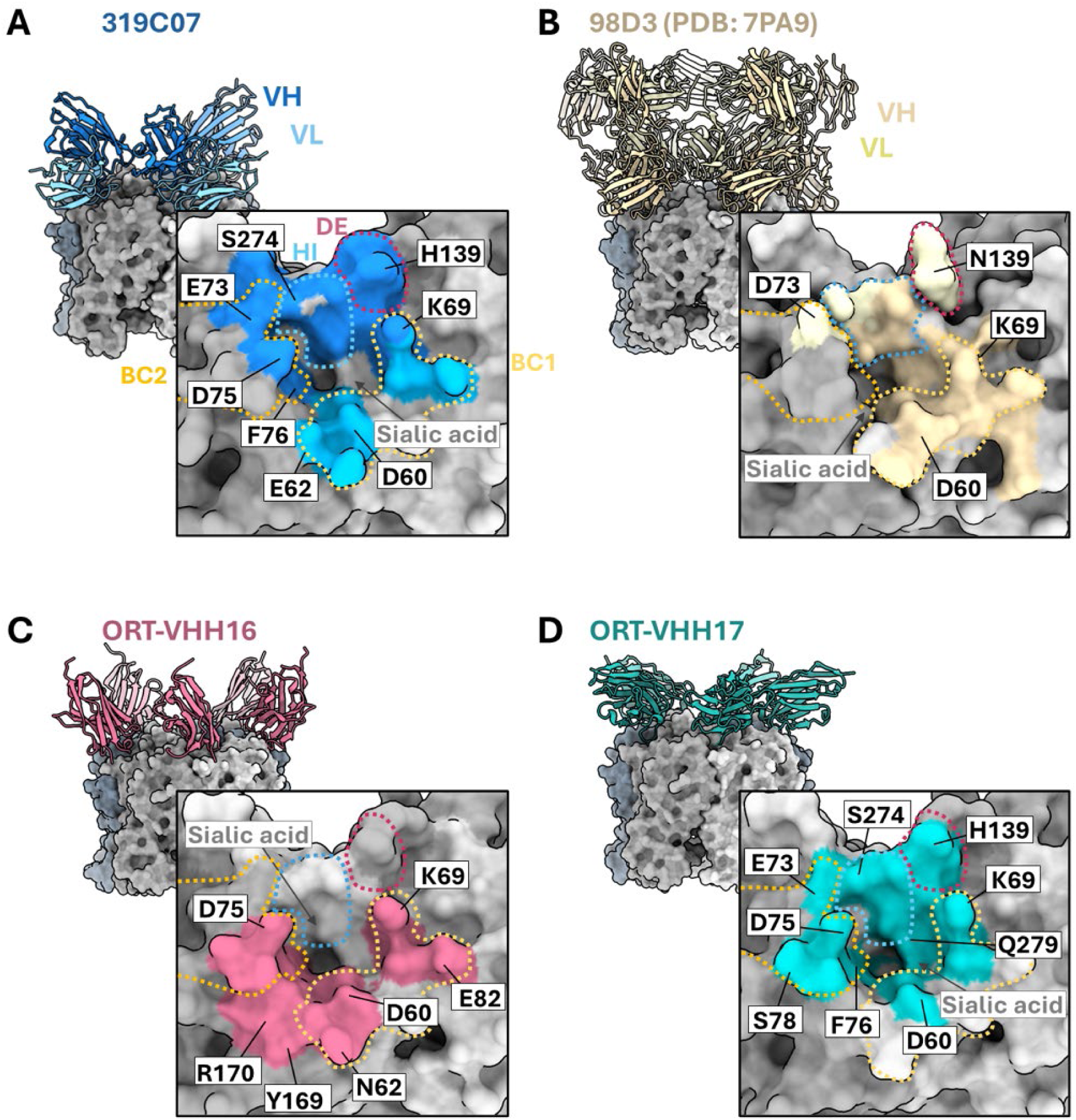
X-ray structures of different mAbs and nanobodies reveal a similar molecular footprint around the sialic acid binding pocket. **A)** X-ray crystal structure of 319C07 scFv in complex with BKPyV VP1. The interacting residues of VP1 with 319C07 are colored dark blue (VH) and light blue (VL). **B)** X-ray crystal structure of Fab 98D3 (PDB <u>7PA9</u>) in complex with JCPyV VP1. The interacting residues of VP1 with 98D3 are colored yellow (HC) and light yellow (LC). **C)** X-ray crystal structure of ORT-VHH16 in complex with BKPyV VP1. The interacting residues of BKPyV VP1 are colored in light magenta. **D)** X-ray crystal structure of ORT-VHH17 in complex with BKPyV VP1. The interacting residues of BKPyV VP1 are colored in cyan. Several of the key interacting residues are annotated.

**Figure 3:**
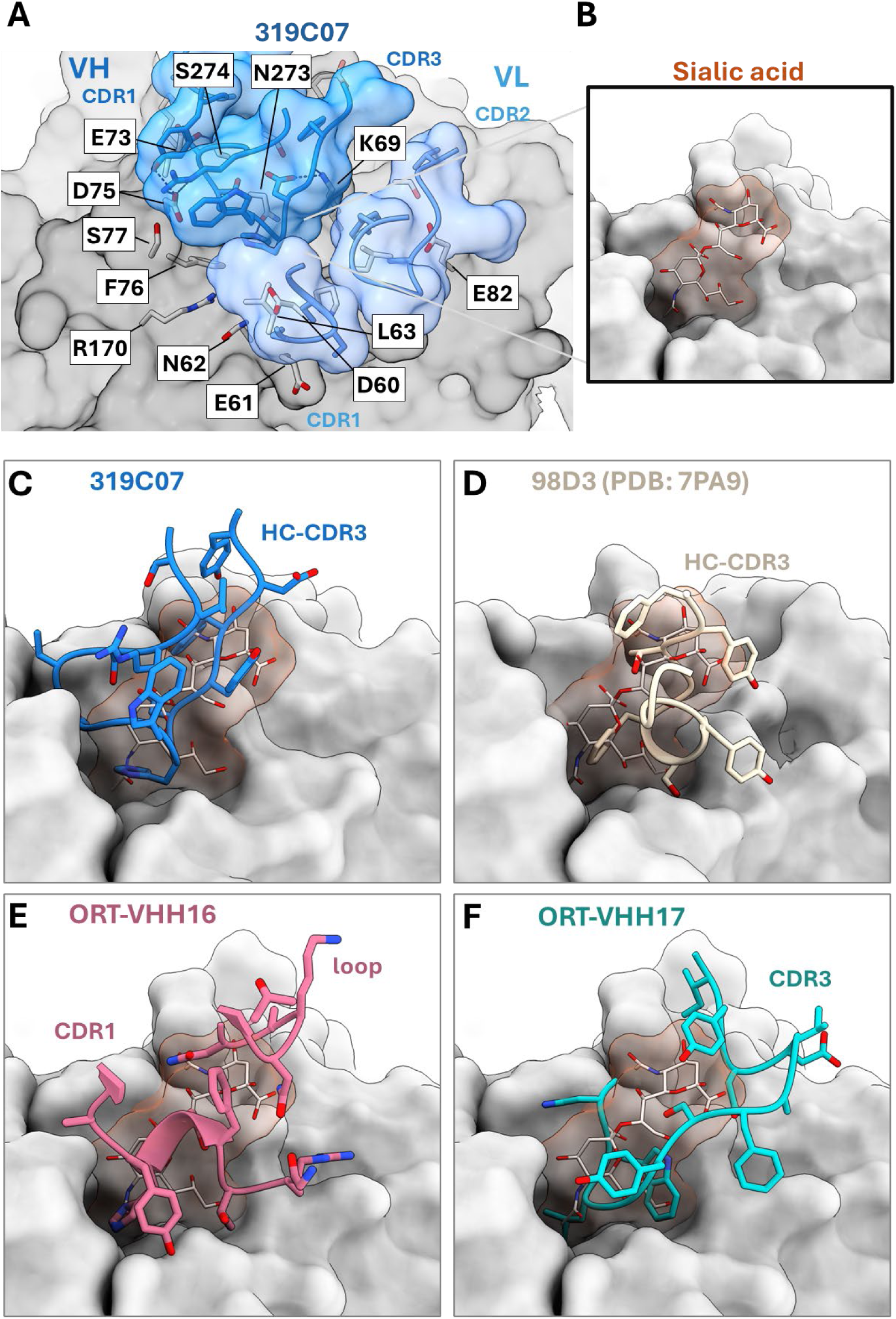
mAbs and nanobodies insert their CDRs into the sialic acid binding pocket. **A)** Closeup of 319C07 scFv interaction with BKPyV VP1. Interacting residues between 319C07 and VP1 are shown. **B-F)** The NeuAc5 glycan (beige with red blob) shown in the VP1 receptor-cleft. CDRs and corresponding residues that occupy the sialic acid binding pocket are shown for 319C07, 98D3, ORT-VHH16 and ORT-VHH17, overlapping the sialic acid molecule (PDB: <u>4MJ0</u>).

The dominant interaction is mediated by the heavy chain CDR-H3 loop, which inserts deeply into the Neu5Ac-binding cleft. The loop forms backbone hydrogen bonds with K69 and additional contacts with E73, F76, and E82, residues implicated in sialylated glycan recognition in glycan-bound VP1 structures. Superposition with Neu5Ac-complexed VP1 (PDB: <u>4MJ0</u>) confirms that the CDR-H3 occupies the same spatial volume as the native receptor, sterically displacing it. By topologically mimicking Neu5Ac, reproducing its spatial positioning and pocket occupancy rather than its precise chemistry, 319C07 sterically occludes receptor binding and directly engages VP1 residues that define the ganglioside-binding site. This structural mimicry likely contributes to the high *in vitro* neutralization potency observed for 319C07 in cell-based assays (**Table 2**). The VL domain of 319C07 provides additional stabilizing interactions. Its CDR-L1, particularly the SSSY motif (residues 27-30), makes polar and van der Waals contacts at the cleft rim, while the CDR-L2 engages peripheral VP1 residues and the CDR-L3 remains structurally disengaged from the cleft.

**Table 2:** *In vitro* neutralization potencies of BKPyV VP1-targeting mAbs, and nanobodies in BKPyV infected human RPTE cell assay

| Compound/mAb/nAb | Description | BPyKV-Dunlop RPTEC Assay |
| --- | --- | --- |
|  |  | EC<sub>50</sub> (±SD) nM |
| 319C07 (Potravitug) | mAb | 0.003 (±0.002) |
| 336F07 | mAb | 0.007 (±0.009) |
| MAU868 | mAb | 0.026 (±0.029) |
| 41F17 | mAb | 0.004 (±0.002) |
| ORT-VHH16* | nAb (single domain heavy chain) | 12.6 (±3.8) |
| ORT-VHH17* | nAb (single domain heavy chain) | 6 (±1.8) |

Notes: EC<sub>50</sub> values (BKPyV DNA by qPCR) are presented as average and standard deviation from at least 3 independent experiments using BKPyV serotype Ia WT VP1 (Dunlop) infected RPTE (human renal proximal tubular) cells at MOI of 0.01. \*VHH16/17 are monovalent, Fc-less heavy chain domains whereas 319C07, 336F07, MAU868 and 41F17 represent bivalent IgG molecules with enhanced avidity and Fc-domains

During the preparation of this manuscript, the structure of full-length Fab-319C07 bound to BKPyV VP1 was reported (PDB: <u>9RM2</u>). Comparison with the scFv complex reveals a nearly identical binding footprint and epitope orientation, indicating that cleft engagement is structurally conserved across antibody formats. A subtle ∼2° rotational shift in the scFv increases BC-loop burial by ∼10 Å² while reducing peripheral contacts, resulting in 3:1 binding per pentamer compared with the 1:1 stoichiometry for the Fab and VP1. This observation confirms the epitope visualized in our structure as format-independent and robust.

Evaluation of the structure of antibody 98D3 bound to JCPyV VP1 (PDB: <u>7PA9</u>), further supports the existence of a receptor-cleft-targeting antibody class. 98D3 is a human monoclonal antibody that showed high affinity JCPyV VP1 binding [24]. Our structural examination showed that 98D3 inserts its HCDR3 loop into the sialic acid binding cleft, bridging the BC-loop and overlapping the native receptor footprint. A phenylalanine residue projects into the hydrophobic pocket typically occupied by the C5 N-acetyl group of Neu5Ac, mimicking the apolar face of the glycan (**Figure 3B**). Although lacking canonical hydrogen bonds, this side chain recapitulates the steric volume and burial characteristics of the native ligand’s C5 moiety, suggesting a comparable mode of receptor interference.

### B. Diverse loop configurations enable convergent glycan-cleft targeting by potent BKPyV-neutralizing nanobodies

To evaluate compact antibody formats with improved access to immunoprivileged compartments such as the renal parenchyma, we selected two potent BKPyV-neutralizing nanobodies for structural analysis. Our high-resolution co-crystal structures of VHH16 and VHH17 (2.6 Å and 2.5 Å, respectively) revealed convergent binding to the sialic acid-binding cleft of VP1, with each paratope occluding key glycan-contacting residues. Despite engaging a markedly smaller epitope than 319C07 (∼700 Å² vs. 915 Å²; ∼77%), and lacking Fc regions and bivalency, both nanobodies exhibited near single digit nanomolar neutralization potency *in vitro* (**Table 2**) suggesting that direct occupation of the receptor-binding cleft may be sufficient to confer potent neutralizing activity. Both nanobodies bind the BC-loop hairpin with 1:1 stoichiometry per VP1 monomer but utilize distinct paratope architectures, approach angles, and loop geometries to efficiently engage the receptor-binding site.

VHH16 achieves cleft occlusion through a rim-spanning configuration dominated by its extended CDR1 (residues G26–Y32), supported by a projecting CDR3 that bridges adjacent VP1 monomers (**Figure 2C**). VHH16 projects its CDR1 residue H31 into the hydrophobic pocket normally occupied by the C5 N-acetyl group of Neu5Ac, where it forms a hydrogen bond with R170, a residue critical for glycan coordination. The insertion of H31 of VHH16 into the cleft is reminiscent of H235 of 319C07, although they adopt differing conformations when accessing the receptor cleft. Additional contacts include a salt bridge between D30 and K69, and hydrogen bonds from Y32 to D60 and N62. The CDR3, centered around Q101, spans across the cleft axis and interacts with both BC1 and BC2 loops of adjacent VP1 monomers, anchoring the paratope over the canyon rim. This architecture sterically caps the receptor-binding site, blocking glycan access through rim-level occlusion (**Figure 3E**).

In contrast, VHH17 inserts a compact CDR3 loop (residues P99-F106) deep into the canyon floor of the sialic acid-binding cleft of VP1, using a hydrophobic triad to anchor within the cleft and a secondary polar network to stabilize the interactions (**Figure 2D**). The indole side chain of its W101 docks into a glove-like hydrophobic cavity formed by VP1 residues L62, L67, P58, and K68. This aromatic insertion mimics the nonpolar face of the Neu5Ac pyranose ring and forms extensive van der Waals contacts across the canyon floor (**Figure 3F**). Flanking residues P99 and V102 form a bilateral hydrophobic clamp that recapitulates the topography of Neu5Ac. P99 aligns spatially with the axial C6 methyl group, while V102 projects its isopropyl side chain into the pocket otherwise occupied by the C5 N-acetyl methyl, engaging with greater surface complementarity than the native ligand or VHH16. N100 bridges these components by forming a stabilizing hydrogen bond with S271 on VP1, while also orienting the loop for optimal insertion. Residues G103-F106 of VHH17 trace the glycan’s natural exit trajectory, projecting over the cleft rim and contacting VP1 residues E77, Q59, and T278. This sterically occludes the exit channel used by extended sialylated glycans, enforcing complete blockade of multivalent glycan engagement. Additional stabilizing interactions are provided by the CDR2 residue R53, which forms a polar contact with D59 of the BC-loop, further anchoring this nanobody. Collectively, VHH17 functions as a rigid, high-affinity Neu5Ac mimetic, replacing the flexible, low-affinity glycan with a preorganized CDR3 loop that reproduces both the spatial footprint and chemical features of the native ligand.

These topological differences between the two nanobodies represent distinct paratope solutions to the steric and electrostatic constraints of the VP1 cleft surface, revealing that the sialic-acid pocket on VP1 can support structurally orthogonal modes of antibody access. Together, the structures of VHH16 and VHH17, alongside those of 319C07 and 98D3, reveal a recurrent focus on the VP1 receptor-binding surface, prompting a classification of antibody epitopes to better define the structural and mechanistic implications of their binding modes.

### **C.** Structural classification of antibody binding modes reveals a shared focus on the BC-loop of BKPyV VP1

To organize the structural diversity of BKPyV-directed antibodies, we systematically analyzed available VP1-antibody complexes in the Protein Data Bank. This enabled classification based on antibody orientation and epitope footprint, revealing two dominant binding configurations. To contextualize these configurations, we used the glycan-bound VP1 structure (PDB: <u>4MJ0</u>) to define the geometry of the sialic acid-binding cleft. This receptor-binding site is shaped primarily by the BC-loop (residues 55-85) and the HI-loop (residues 270-277), which form the floor and lateral edges of the cleft, respectively (**Figure 1**). The DE-loop (132-142) contributes minimally to the receptor interface. Within the cleft, polar contacts from the HI-loop’s N273-T277 segment and the apex of the BC-loop coordinate the terminal Neu5Ac moiety of host gangliosides. This configuration provides a structural reference point for mapping antibody interactions that may sterically block, mimic, or compete with receptor engagement.

Apical binding antibodies engage the sialic acid-binding cleft of VP1 through direct insertion or rim-level occlusion, often mimicking the receptor or blocking its access sterically. This class includes 319C07, 98D3, VHH16, and VHH17, all of which either bury their paratopes into the Neu5Ac-binding pocket or occlude it while making extensive contacts with the underlying BC-loop, and in most cases, the adjacent HI-loop residues **(Figure 4A).**

**Figure 4:**
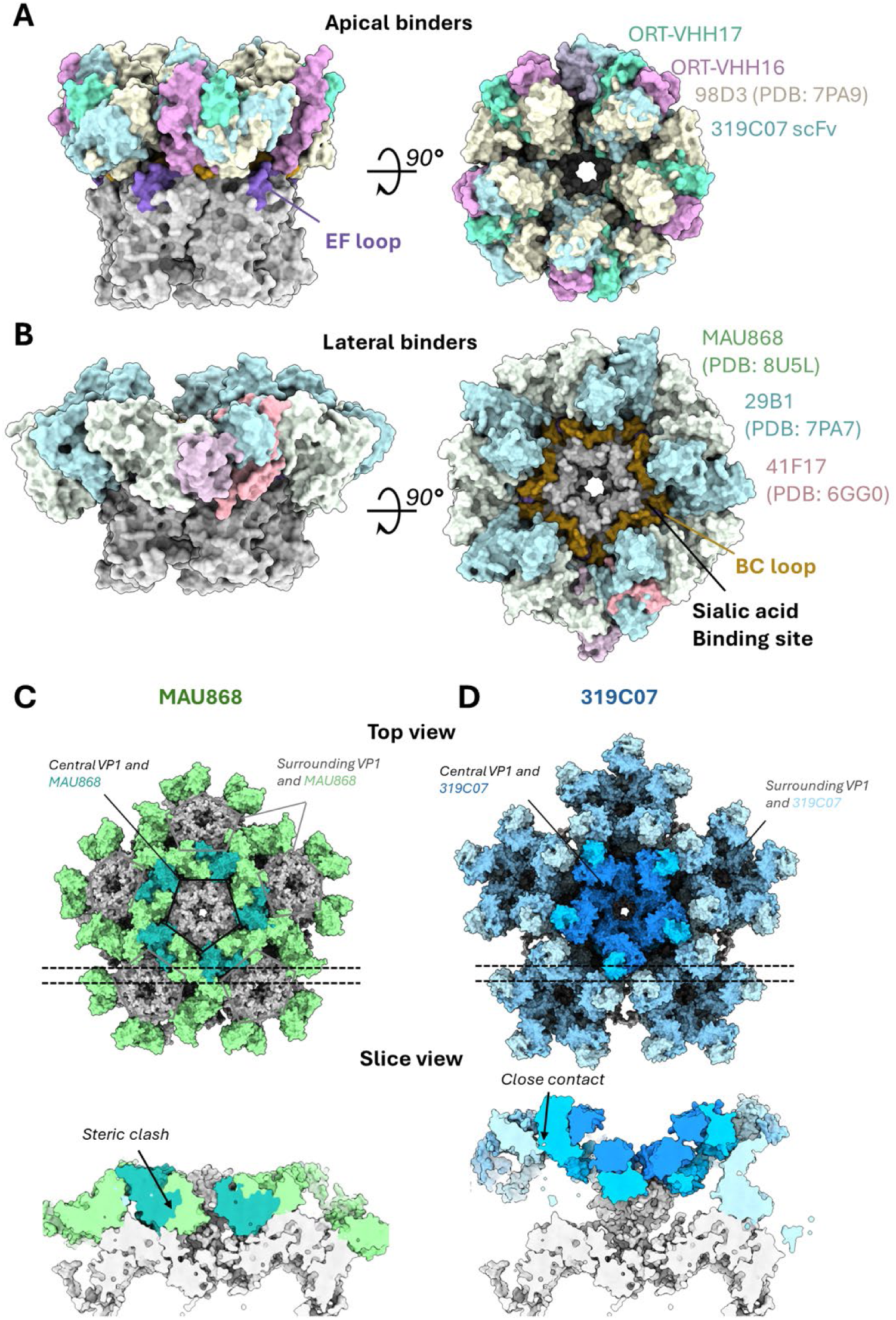
Antibodies targeting BKPyV VP1 can be classified into two distinct groups. Surface representation of mAbs and nanobodies that bind apically to the sialic acid binding pocket colored yellow (98D3), green (ORT-VHH17), pink (ORT-VHH16) and blue (319C07) (**panel A**), and laterally, colored light green (MAU868), cyan (29B1) and pink (41F17) (**panel B**). BKPyV VP1is colored grey with the BC and EF loops colored gold and purple, respectively. Surface representation of clinical stage antibodies modeled on the pentavalent face of an assembled BKPyV (PDB: 6GG0) to illustrate steric constraints of lateral binders like MAU868 (**panel C and D**).

Lateral binders approach the capsid radially from an oblique or peripheral angle, engaging surfaces that lie outside the sialic-acid cleft. MAU868 and 41F17 exemplify this class. MAU868 binds primarily along the EF loop ridge with limited BC-loop interaction, while 41F17 requires fully assembled particles to engage an inter-pentamer epitope formed by adjacent VP1 units, indicating a dependence on intact capsid geometry and symmetry (**Figure 4B)**. One lateral binder of particular interest is 29B1 (PDB: <u>7PA7</u>), which occupies a distinct epitope niche.

Although it lacks direct contact with the Neu5Ac-binding site, it engages the BC-loop extensively from a downward-facing, laterally oblique angle. Its paratope spans cleft-adjacent and lateral surfaces, including portions of the EF loop, forming a hybrid footprint anchored on the BC-loop. This mode highlights the capacity of lateral binding antibodies to simultaneously access BC-loop elements, reinforcing the immunodominant character of the BC-loop.

These structural comparisons reveal that the BC-loop serves as a unifying axis of VP1 antibody recognition across distinct classes. Despite divergent approach angles and neutralization mechanisms, this shared epitope dependence highlights the BC-loop as the immunodominant, and structurally accessible surface most consistently targeted by antibodies. Its prominence across binding classes and formats including mAbs, nanobodies, and naturally elicited antibodies underscores its immunodominance and structural accessibility. Notably, therapeutic antibodies from both major classes, 319C07 (apical) and MAU868 (lateral), have advanced into clinical evaluation, underscoring the translational relevance of binding orientation and epitope architecture [15, 16].

Taken together, these observations frame the BC-loop not only as the structural epicenter of antibody recognition and a frequent target of the humoral response, but also as a focal point of immune pressure. Its repeated targeting raises critical questions about its mutational tolerance, the emergence of escape variants, and its suitability as a stable epitope for the design of broadly effective therapeutics.

### **D.** Comprehensive mining of public BKPyV VP1 sequences reveals extensive natural variation at the BC-loop

The striking convergence of multiple neutralizing antibodies on the sialic acid-binding cleft and the BC-loop, led us to determine whether this region is conserved across BKPyV isolates, or if its immunodominance comes at the cost of genetic instability and viral fitness. To generate a comprehensive, population-level map of VP1 diversity with clinical relevance, we curated all available BK polyomavirus protein sequences from GenBank (NCBI), yielding 3,048 entries.

After stringent quality filtering and de-duplication, we identified 923 full-length VP1 sequences corresponding to 923 unique genome accessions. From these, we derived a final set of 189 non-redundant, full-length sequences spanning genotypes I-IV, which served as the primary dataset for our entropy, conservation, and mutational burden analyses. Inclusion of partial sequences (353 non-redundant entries) in cross-validation analyses confirmed the robustness of our conclusions. These sequences represent all four major genotypes and their subtypes, reflecting the unweighted sampling distribution of BKPyV in public clinical and research studies; genotypes I and IV were most abundant, consistent with global prevalence.

Shannon entropy analysis of the 189 BKPyV VP1 sequences revealed an asymmetric landscape of amino acid variability, sharply concentrated within the BC-loop (residues 55-85) (**Figure 5A**). Entropy values at positions E61, K69, D77, and S79 exceeded 1.0 bits, indicating multiple co-dominant residues at these positions and suggesting localized immune-driven diversification.

**Figure 5:**
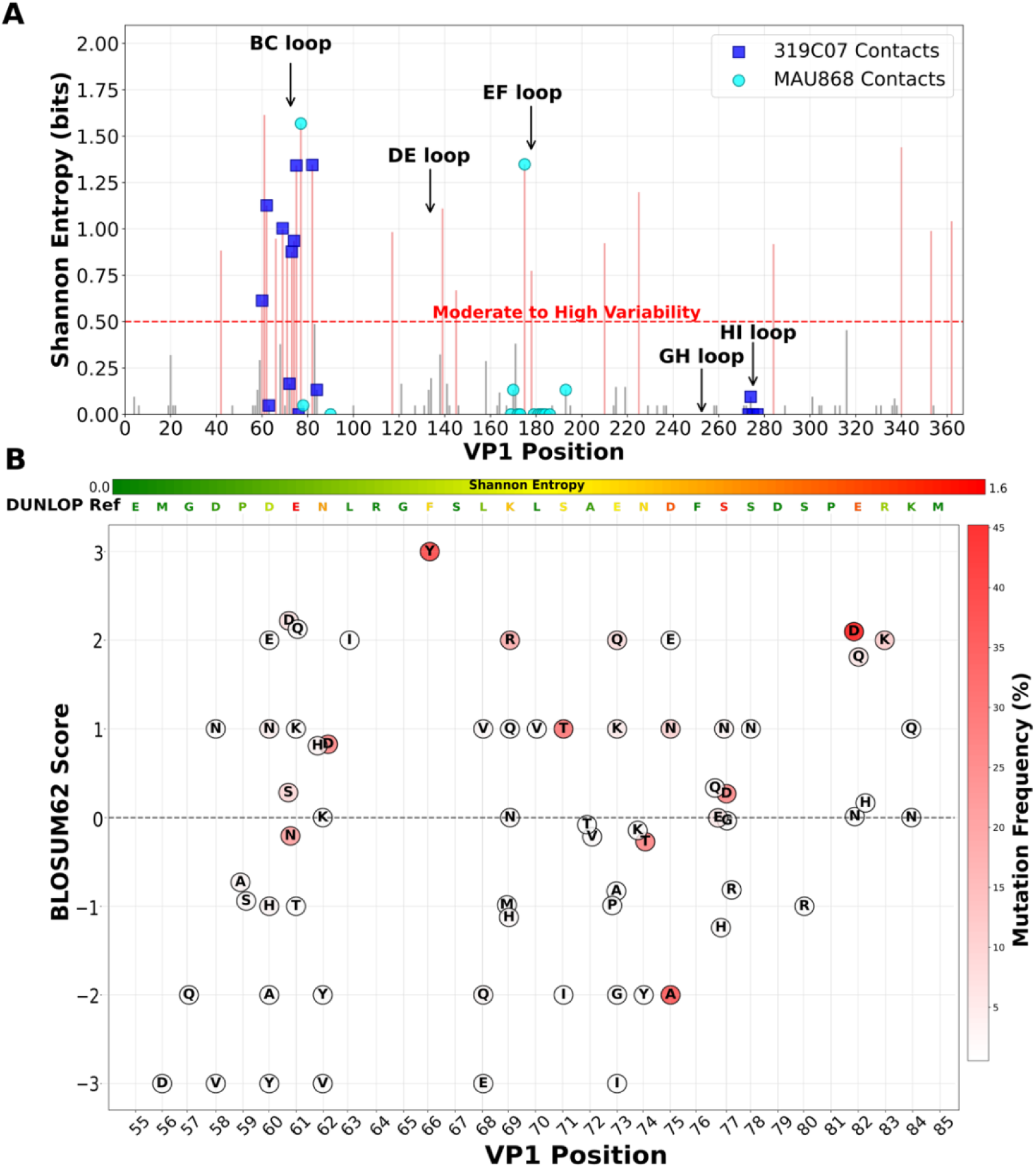
A) Shannon entropy analysis of BK polyomavirus VP1 protein reveals variable regions coinciding with antibody binding sites. Shannon entropy values (bits) are plotted for each amino acid position in the VP1 protein sequence. Bars are colored grey for positions with low variability (≤0.5 bits) and red for positions with moderate to high variability (>0.5 bits), as indicated by the dotted red threshold line. Major structural loops are annotated: BC loop (positions 55-85), DE loop (positions 132-142), EF loop (positions 167-181), GH loop (positions 245-257), and HI loop (positions 270-277). Antibody contact residues are overlaid as blue squares for monoclonal antibody 319C07 and cyan circles for MAU868, demonstrating that several high-entropy positions correspond to antibody binding sites. The analysis reveals that antigenic regions, particularly within the BC and EF loops, exhibit elevated sequence variability, suggesting these sites are under immune selection pressure. **B) Scatter plot of amino acid substitutions in the VP1 BC-loop region (positions 55-85).** Each circle represents a mutation with color intensity indicating frequency (white- to-red scale, right color bar) and black letters showing the substituted amino acid. BLOSUM62 scores (y-axis) quantify the physicochemical impact of substitutions, with positive values indicating conservative changes and negative values indicating radical changes (dashed line at y=0). The Dunlop reference sequence (top axis) is color-coded by Shannon entropy to show evolutionary conservation (green = conserved, yellow = moderate, red = variable; bottom color bar). Points are jittered to prevent overlap while maintaining positional accuracy.

Notably, BC-loop conservation varies dramatically across BKV serotypes, with ST1 showing 36.9% of isolates maintaining perfect conservation (0 mutations), while ST3 exhibits the most extensive remodeling, with only 12.5% conserved isolates. The remainder show 1-11 substitutions across serotypes, with mean mutation burdens ranging from 1.29 (ST2) to 7.12 (ST3), underscoring serotype-specific patterns of mutational remodeling in this critical antibody-binding region. Importantly, the BC-loop accounts for a disproportionate 27.8-53.3% of total VP1 mutations across serotypes, despite representing only ∼10% of the protein sequence. The EF loop (residues 167-181) emerged as the next most variable region, though with overall lower entropy than the BC-loop and contributes to the lateral epitope surfaces bound by MAU868 and 41F17. In contrast, the DE, GH, and HI-loops showed lower but non-negligible variability, while the β-barrel core remained uniformly conserved.

To evaluate the biochemical permissiveness of substitutions within these loops, we mapped all observed variants to BLOSUM62 scores, which estimate the evolutionary likelihood of amino acid replacements based on physicochemical similarity. Variants such as E61N (19.5%), N74T (24.7%), D75A (33.7%), and S79D (23.7%) showed negative BLOSUM62 scores and substantial shifts in charge or polarity, consistent with immune selection at structurally flexible positions (**Figure 5B**). ConSurf projection of per-residue conservation onto the VP1 pentamer surface reinforced this pattern of variable residues clustered predominantly at the apical surface of BKPyV VP1, especially within the BC-loop, while structurally buried residues and inter-pentamer interfaces remained highly conserved (**Figure 6**). Notably, residues within the footprints of 319C07, VHH16, VHH17, MAU868, and 41F17 aligned with entropy-enriched surface patches, with the greatest overlap observed for sialic acid cleft-binding antibodies such as 319C07 and VHH16/17, reinforcing that these frequent and biophysically disruptive mutations directly impact therapeutically targeted surfaces.

**Figure 6:**
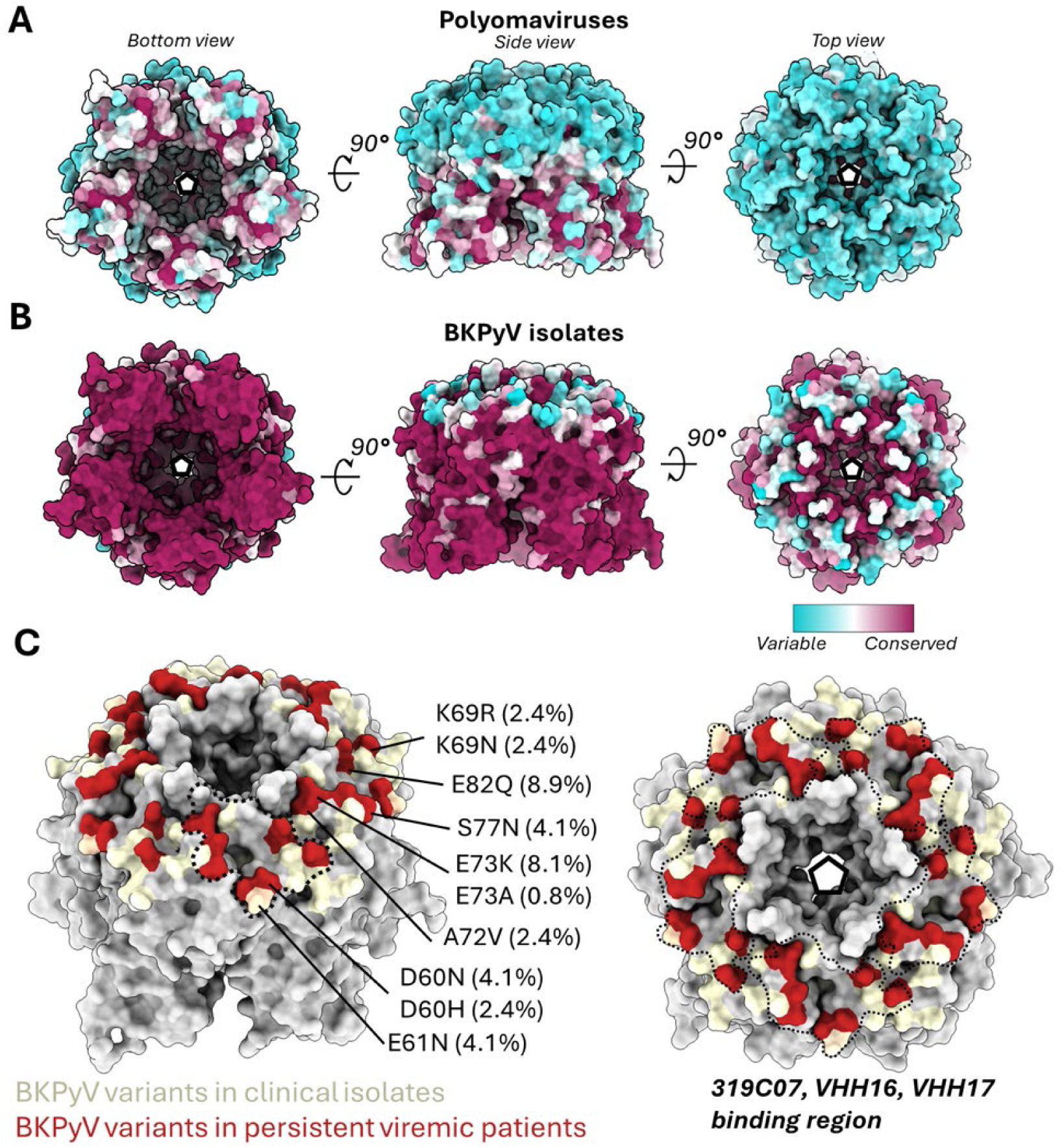
Immune-exposed regions of the polyomavirus VP1 capsids are highly variable. Conservation analysis of VP1 using sequences across the polyomaviridae family (**panel A**) and across all BKPyV VP1 clinical isolates (**panel B**). The surface of BKPyV VP1 is colored according to conservation with cyan representing variable, while maroon represents conserved residues. Across both sequences the BC-loops are especially variable. **C)** Residues that are mutated in the BKPyV VP1 BC-loop for clinical isolates are shown in yellow while mutations associated with confirmed persistent viremia patients are shown in red and further annotated with their prevalence. The figure is a collective representative of viral variants from multiple studies.

Serotype-resolved sequence alignment revealed several additional layers of epitope variability. Multiple BC-loop residues, such as E61, N62, E73, and D77, exhibited independent variation across genotypes, consistent with convergent immune selection. Several isolates harbored compound substitutions at adjacent epitope residues, suggesting structural remodeling beyond single-point escape. Furthermore, low frequency but biochemically disruptive substitutions, including E61K and S79D, highlighted the biochemical breadth of tolerated variation. Even within the conserved HI-loop, S276R/K from the NSSGT motif showed natural mutations in public sequences from genotypes II and IV, revealing a frequency of 7.1% and 2.3% for R and K respectively. Together these patterns collectively reveal the challenges associated with single-epitope or pan-serotype antibody strategies directed at the BC-loop, positioning it at the mutational epicenter of immune pressure and VP1 evolution.

### **E.** Patient-derived VP1 variants from viremic patients reveal recurrent mutations in antibody-contact residues of the BC-Loop

To examine whether the observed BC-loop variability bears clinical consequence, we surveyed published VP1 sequences from transplant recipients with persistent viremia and integrated data from public isolates with known clinical annotations [22, 25–30]. Studies from multiple independent transplant cohorts, report longitudinal or cross-sectional sampling from patients with sustained high-level viremia, delayed viral clearance, or biopsy-proven polyomavirus-associated nephropathy (PVAN). These cohorts consistently describe the emergence of VP1 mutations during persistent viremia, particularly in patients classified as non-controllers (individuals with sustained high-level viremia, typically >10⁴ copies/mL).

Across studies, we observed a striking convergence of substitutions within the BC-loop in non-controlling patients, highlighting residues we independently identified as entropically enriched and mutationally diverse in our broader population-scale analysis. These included high-frequency substitutions such as K69R/N/Q, E72V, E73K/A/Q, S77N, E82Q. In longitudinally sampled patients, VP1 variants accumulated stepwise, often appearing in combination, suggesting compound escape (**Table 3**). Notably, several map directly within the epitope footprints of therapeutic antibodies including 319C07 and VHH16/17. For example, K69, a central contact for all three, was frequently substituted, and the NEA27 isolate (from a PVAN patient) harbors E62H and K69H, two putatively disruptive mutations directly within the 319C07 interface.

**Table 3:** Natural prevalence of variants BKPyV VP1 sequences in clinical samples derived from the GenBank database.

| BKPyV<br>VP1 AA<br>position | ST1a<br>WT Ref<br>AA | ST1 | ST2 | ST3 | ST4 |
| --- | --- | --- | --- | --- | --- |
| 60 | D | D:91.1% H:2.4% N:4.1%<br>E:2.4% | D:100% | D:87.5% A:12.5% | D:88.6% H:4.5%<br>N:4.5% Y:2.3% |
| 61 | E | E:87.8% N:8.1% D:0.8%<br>K:1.6% Q:1.6% | E:21.4% D:64.3%<br>N:14.3% | E:62.5% D:25%<br>T:12.5% | N:56.8% E:6.8%<br>S:36.4% |
| 62 | N | N:87.8% D:6.5% H:4.1%<br>Y:0.8% K:0.8% | N:92.9% D:7.1% | N:75.0% H:12.5%<br>V:12.5% | D:84.1% N:15.9% |
| 63 | L | L:100% | L:100% | L:87.5% I:12.5% | L:100% |
| 68 | L | L:95.1% V:3.3% E:1.6% | L:100% | L:50.0% Q:50% | L:100% |
| 69 | K | K:93.5% M:0.8% N:2.4%<br>R:2.4% Q:0.8% | K:100% | K:50% H:50% | R:65.9% K:34.1% |
| 72 | A | A:96.7% V:2.4% T:0.8% | A:100% | A:100% | A:100% |
| 73 | E | E:81.3% Q:8.1% K:8.1%<br>I:0.8% G:0.8% A:0.8% | E:100 % | P:12.5% E:87.5% | E:90.9% Q:6.8%<br>G:2.3% |
| 74 | N | N:91.9% T:5.7% K:1.6%<br>Y:0.8% | N:100% | N:100% | T:90.9% N:9.1% |
| 75 | D | D:80.5% N:13.8% A:4.9%<br>E:0.8% | A:92.9% D:7.1% | D:50.0% A:50.0% | A:90.9% D:9.1% |
| 76 | F | F:100% | F:100% | F:100.0% | F:100% |
| 82 | E | E:70.7% D:19.5% N:0.8%<br>Q:8.9% | D:92.9% H:7.1% | E:37.5% D:62.5% | D:100% |
| 84 | K | K:97.6% N:0.8% Q:1.6% | K:100% | K:100% | K:100% |
| 273 | N | N:100% | N:100% | N:100% | N:100% |
| 274 | S | S:100% | S:92.9% R:7.1% | S:100% | S:97.7% K:2.3% |
| 275 | S | S:100% | S:100% | S:100% | S:100% |
| 277 | T | T:100% | T:100% | T:100% | T:100% |
| Notes: AA = amino acid. ST1a = serotype 1a, ST2 = serotype 2, ST3 = serotype 3 and ST4 = serotype 4; WT = wildtype |  |  |  |  |  |

These substitutions exhibited distinct evolutionary trajectories across diverse patient backgrounds, with serotype-specific mutation burdens revealing differential immune selection pressures. ST1 and ST2 show relatively conservative patterns (means 1.55 and 1.29 mutations, respectively), while ST4 demonstrates intermediate remodeling (mean 2.02). In contrast, ST3 exhibits extreme mutational burden (mean 7.12) with a striking bimodal distribution, 75% of ST3 sequences harbor ≥5 BC-loop mutations compared to only 7.1-11.4% in other serotypes. This concentrated mutational remodeling, particularly evident in ST3’s extensive amino acid substitutions, indicates a region under intense immune selection that tolerates substantial biochemical variation while maintaining viral fitness and infectivity. To determine whether these clinical mutations reflect stochastic drift or selective pressure, we cross-referenced them against entropy and BLOSUM62 scores in our curated dataset of 189 non-redundant VP1 sequences.

Many patient-derived substitutions, including E61N (19.5%) and S77D (23.7%), occurred at sites with high entropy and low substitution similarity, often involving charge-altering or polarity-shifting mutations, supporting a model of antibody driven immune escape.

These findings provide a translationally relevant correlation to the widespread mutational tolerance and structural accessibility described in our larger population-scale bioinformatics analysis of BC-loop variability in BKPyV VP1. They define the BC-loop as an active site of immune-driven diversity *in vivo*, particularly in patients failing to control viral replication. The overlap between real-world clinical mutations and neutralizing antibody footprints emphasizes the vulnerability of targeting the BC-loop (**Figure 6**). These observations support a model in which persistent viremia provides the substrate for potential therapeutic non-responsiveness of BC-loop targeting antibodies and expose two converging liabilities of the BC-loop: its role as a site of preexisting serologic non-responsiveness in patients with uncontrolled viremia, and its capacity to support immune escape through biochemically disruptive yet functionally tolerated substitutions.

This pattern of variation observed in patients with persistent BK viremia, mirrors therapeutic escape trajectories seen in structurally homologous polyomavirus capsids. In a murine polyomavirus model of JCPyV infection, monoclonal antibody treatment selected mutations within the apical BC and HI-loops, orthologous to BKPyV’s receptor-binding-cleft epitope, resulting in therapeutic antibody resistance, demonstrating that this surface supports antigenic plasticity under immune pressure [31].

### **F.** Patient-derived VP1 BC-loop substitutions abrogate binding of sialic acid cleft-targeting antibodies

To assess the impact of clinically observed VP1 variation on therapeutic antibody binding, we performed biolayer interferometry (BLI) against a panel of recombinant VP1 pentamers spanning all four BKPyV serotypes, reference subtypes, and patient-derived variants enriched in persistent viremia (**Table 4**). This panel included both genotype-specific reference sequences (e.g., Dunlop 1a, M23122 III-AS, NEA-27 III) and variants carrying BC-loop and HI-loop substitutions documented in viremic transplant recipients, including “non-controller” patients with sustained high viral loads [22, 25–27].

**Table 4:** Clinically relevant VP1 sequences used in this study

| Accession # | Type | Reference | Clinical Relevance/Notes |
| --- | --- | --- | --- |
| KP412983.1 (Cont) | 1a | Henriksen 2015 | Serotype 1a Reference (Dunlop)* |
| KT896261.1 | | Domingo-Calap et al 2024 | Urine (Viral load $1.16 \times 10^8$ copies/mL) |
| LR216160.1 |  | Van DT 2019 | Blood: collected from an immunosuppressed patient |
| MW587977.1 | | Addetia, A., et al 2021 | Urine: from immunosuppressed patient (Viral load $3.90 \times 10^7$ copies/mL) |
| OQ230860.1 | 1b-1 | Mineeva-Sangwo et al 2023 | Blood: Serotype 1b-1 Reference; KTx patient; Immunosuppressed |
| LR216078.1 |  | Van DT 2019 | Urine: collected from an immunosuppressed patient |
| LR216092.1 |  | Van DT 2019 | Urine: collected from an immunosuppressed patient |
| LR216094.1 |  | Van DT 2019 | Urine: collected from an immunosuppressed patient |
| LR216105.1 |  | Van DT 2019 | Urine: collected from an immunosuppressed patient |
| OQ230837.1 |  | Mineeva-Sangwo et al 2023 | Urine: KTx patient with persistent viremia (patient 3; 2nd timepoint) |
| OQ230844.1 | 1b-2 | Mineeva-Sangwo et al 2023 | Blood: Serotype 1b-2 Reference; KTx patient; Immunosuppressed |
| KT896383.1 | | Domingo-Calap et al 2024 | Urine: (Viral Load $4.65 \times 10^7$ copies/mL); KTx patient |
| OQ230849.1 |  | Mineeva-Sangwo et al 2023 | Urine: KTx patient with persistent viremia (patient 3; 2nd timepoint) |
| 1b2 + RQN |  | McIlroy (2020) | Urine: persistent viremic "non-controller" patients |
| 1b2 + NNVQ |  | McIlroy (2020) | Urine: persistent viremic "non-controller" patients |
| LR216239 (Cont) | 1c | Van DT 2019 | Urine collected from an immunosuppressed patient |
| OM938900.1 (Cont) | II | Sangwo.....Snoeck 2023 | Serotype II Reference |
| OQ230840.1 |  | Sangwo.....Snoeck 2023 | Urine: KTx patient with persistent viremia (Patient 6; 2<sup>nd</sup> Timepoint) |
| KT896240.1 | | Domingo-Calap et al 2024 | Urine (Viral Load $4.87 \times 10^5$ copies/mL); KTx patient |
| KT896285.1 | | Domingo-Calap et al 2024 | Urine (Viral Load $1.49 \times 10^9$ copies/mL); KTx patient |
| M23122_AS (Cont) | III | Tavis et al 1989 | Serotype III Reference |
| KF055891_KT514USA |  | Kapusinszky et al 2013 | Blood: Pediatric Renal Transplant Patient with Nephropathy |
| AB365139_NEA-27 (Cont) |  | Yogo et al 2008 | Serotype III Reference |
| LR216116.1 (Cont) | IVa-1 | Van DT 2019 | Urine: collected from an immunosuppressed patient |
| LR216102.1 |  | Van DT 2019 | Urine collected from an immunosuppressed patient |
| AB365147.1 (Cont) | IVb-1/2 | Yogo et al 2008 | Serotype IVb-1,2 Reference |
| LR216133.1(Cont) | IVc-1 | Van DT 2019 | Urine collected from an immunosuppressed patient |
| LR216137.1 |  | Van DT 2019 | Urine collected from an immunosuppressed patient |
| OQ230846.1 (Cont) | IVc-2 | Sangwo.....Snoeck 2023 | Serotype IVc-2 Reference |
| OQ230836.1 |  | Sangwo.....Snoeck 2023 | Urine: KTx recipients with persistent viremia (patient 2<sup>cdv</sup>; 2nd timepoint |
| IVc-2 + SKAN |  | McIlroy (2020) | Urine: persistent viremic "non-controller" patients |

Notes: KTx = kidney transplant; cdv = cidofovir;

We evaluated five antibodies, the cleft-invading 319C07, nanobodies VHH16 and VHH17, and the lateral binding MAU868, the latter included as a control antibody for benchmarking due to its minimal dependence on the BC-loop (**Figure 4**). IgG 336F07 served as an internal comparator from the same screening lineage as 319C07 (<u>WO 2021/250097</u>), though its epitope remains structurally undefined.

Nanobodies VHH16 and VHH17, despite nanomolar potency against canonical VP1 (Dunlop 1a), lost binding to most clinical variants with BC-loop substitutions. Binding loss was most pronounced for K69N and E73K, both individually sufficient to abolish measurable interaction (NB, red) (**Figure 7A**). These results reflect the narrow tolerance of their compact paratopes and strict dependence on the receptor-binding cleft.

**Figure 7:**
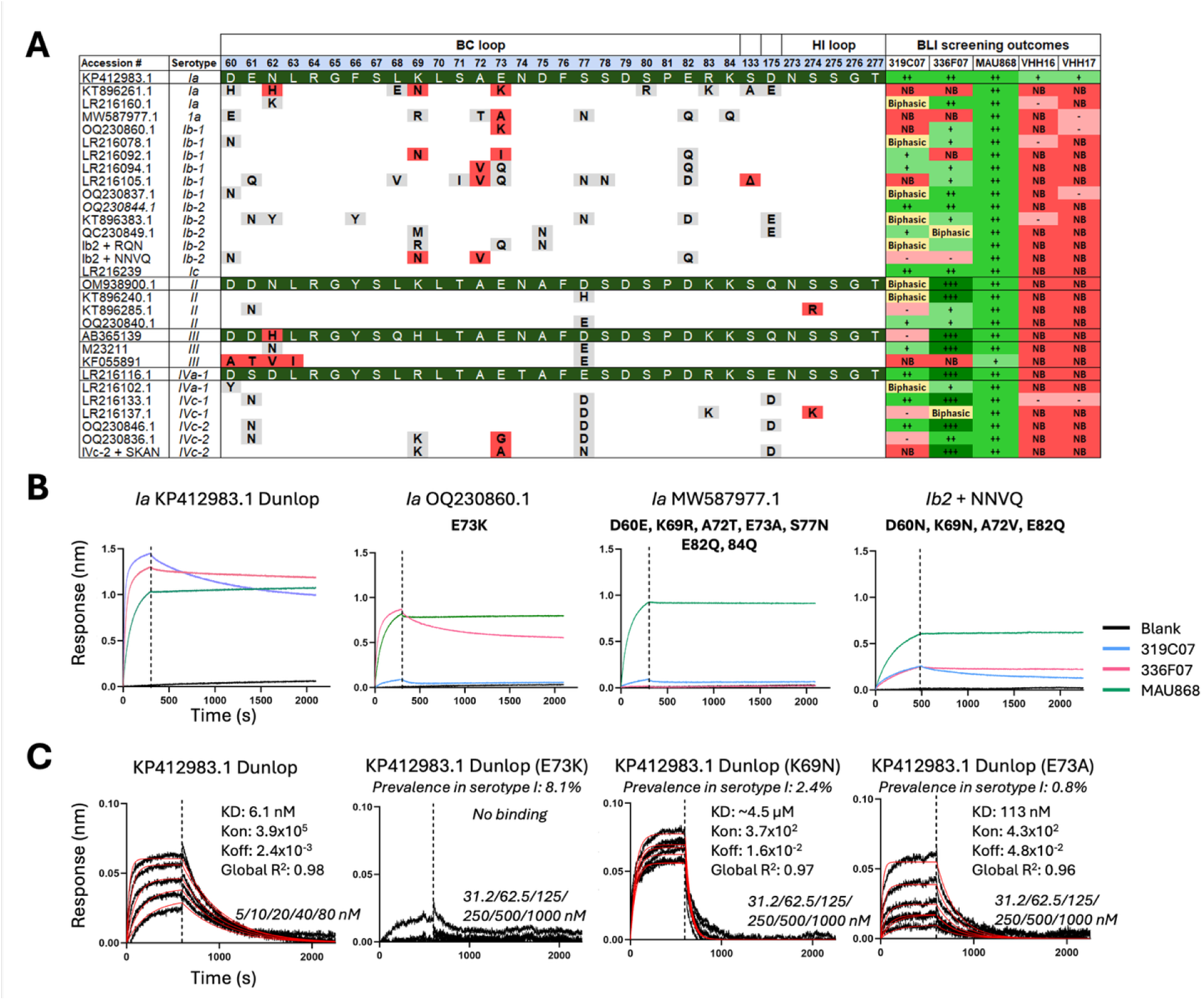
BLI studies to determine binding affinities of clinically relevant VP1 sequences. **A)** Binding responses were classified using a graded intensity scale: dark green (+++, ++) indicated full binding, pale green (+) partial retention, yellow ("Biphasic") suggested altered kinetics, and red ("NB") indicated complete loss of measurable interaction. **B)** Representative BLI sensorgrams of mAbs evaluated in this study **C)** Representative sensorgrams of single point mutations in Dunlop background. 319C07 was immobilized and a concentrations gradient of BKPyV VP1 was tested (concentrations used are indicated on each graph) as described in the Materials and Methods. The raw data (black) was fitted (red) using Prism (Graphpad).

In contrast, 319C07 retained measurable binding to a broader subset of VP1 variants, including pentamers from multiple genotypes, including Ia (Dunlop), Ib2, II, III-AS (M23122), and IVa-1. However, its binding was consistently attenuated by naturally occurring BC-loop substitutions. Complete loss of binding (NB; dark red) was observed against isolates encoding E73K, K69N, or the persistent viremic "non-controller" patients (McIlroy Quad IVc-2 variant), which combines multiple BC- and HI-loop substitutions. Variants such as the VQQ mutant (A72V + E73Q), S274R, and additional viremic patient isolates exhibited either reduced maximal binding (B_max) or biphasic association profiles, consistent with altered loop conformation or asymmetric epitope presentation and differential engagement. As expected, MAU868 retained binding (+/++, green) across all VP1 variants, including those that completely abrogated 319C07 and nanobody interaction. This confirms that the observed losses are specific to sialic acid-cleft- targeting modalities, and not a generalized consequence of VP1 misfolding or an experimental artifact.

To isolate the contribution of individual residues, we generated single-point VP1 mutants, introducing K69N, E73A, and E73K into the isogenic Dunlop background (**Figure 7C**). The binding deficits were recapitulated, showing K69N substitution reduced 319C07 affinity by >3 orders of magnitude (K_D_ ≈ 4.5 µM) and E73K consistently abolished binding entirely. E73A preserved 319C07 affinity but lowered B_max indicating partial occlusion or conformational rearrangement of the epitope. These effects align with prior structural work showing that substitutions at A72 and E73 can remodel the BC-loop disrupting this epitope by inducing a BC2 loop flip, altering the orientation of residues 73-75 in monomers (VQQ) or a subset (E73A) [32]. Although these residues lie outside the Neu5Ac-contacting BC1 floor, such rearrangements remodel the cleft-adjacent epitope landscape. Antibodies like 319C07, which span the BC apex, may therefore encounter a heterogeneous epitope, producing biphasic binding kinetics. In contrast, VHH16 and VHH17, which do not contact residues 71-75, are unlikely to be affected by BC2 loop flipping, consistent with the absence of biphasic association profiles for these nanobodies.

Together, these findings expose the sensitivity of sialic acid-binding-cleft-directed antibodies to naturally occurring sequence variation within the BC and HI-loops of VP1. Our binding data defines a ceiling on the mutational tolerance of sialic acid-cleft-targeting antibodies, even those with optimized binding geometries. While 319C07 retained partial breadth relative to nanobodies, both were ultimately constrained by the inherent plasticity of exposed surface loops. Importantly, many of the disruptive mutations identified in this screen, including K69N and E73K, were previously found to be enriched in VP1 sequences from patients with persistent viremia [22, 25–27]. Several of these mutations occur at high-entropy, low BLOSUM62 score positions in our population dataset, indicating evolutionary plasticity and physicochemical divergence from reference residues.

These data confirm that BC-loop mutations frequently encountered in viremic transplant recipients confer selective resistance to individual neutralizing antibodies. This convergence between preexisting viral escape pathways and experimental binding deficits highlights the immunologic liability of targeting the BC-loop.

### **H.** Patient-derived VP1 BC-loop variants drive 319C07 escape

To assess the impact of the affinity loss of VP1 BC-loop variants observed in our BLI studies, we tested a select set of variants using a pseudovirus (PsV) reporter assay system described previously [33]. BKPyV WT or VP1 variant sequences were introduced into pIaw vector and co-transfected with plasmids carrying VP2, VP3 and a nano-luciferase reporter plasmid to produce pseudovirions as detailed in material and methods. To test the neutralization potency of mAbs, PsVs were preincubated with a serial dilution of 319C07 and added to HEK293TT cells. An IgG isotype and cidofovir served as controls. Following 3 days of incubation nano-luciferase was measured. The dose responses curves for antibody and cidofovir are shown in **Figure 8A-C**. PsV carrying a VP1 E73K substitution in the BC-loop (OQ230860.1) resulted in a profound >50,000- fold loss of neutralizing potency of 319C07 (**Figure 8D**) consistent with no binding observed in BLI studies (**Figure 7**). Similar results were observed for VP1 from MW587977.1 (1a D60E, K69R, A72T, E73A, S77N, E82Q, E84Q) and LR216137.1 (IVc-1 E77D, R83K and S274K) while the VP1 from OQ230836.1 (IVc-2 S61N, R69K, E73G, S77G) showed a higher EC_50_ value of 0.1 nM, which in comparison to the 1a WT represents a 500-fold lower neutralizing potency. As expected, the IgG isotype control and cidofovir showed no measurable neutralization up to the highest concentration tested.

**Figure 8:**
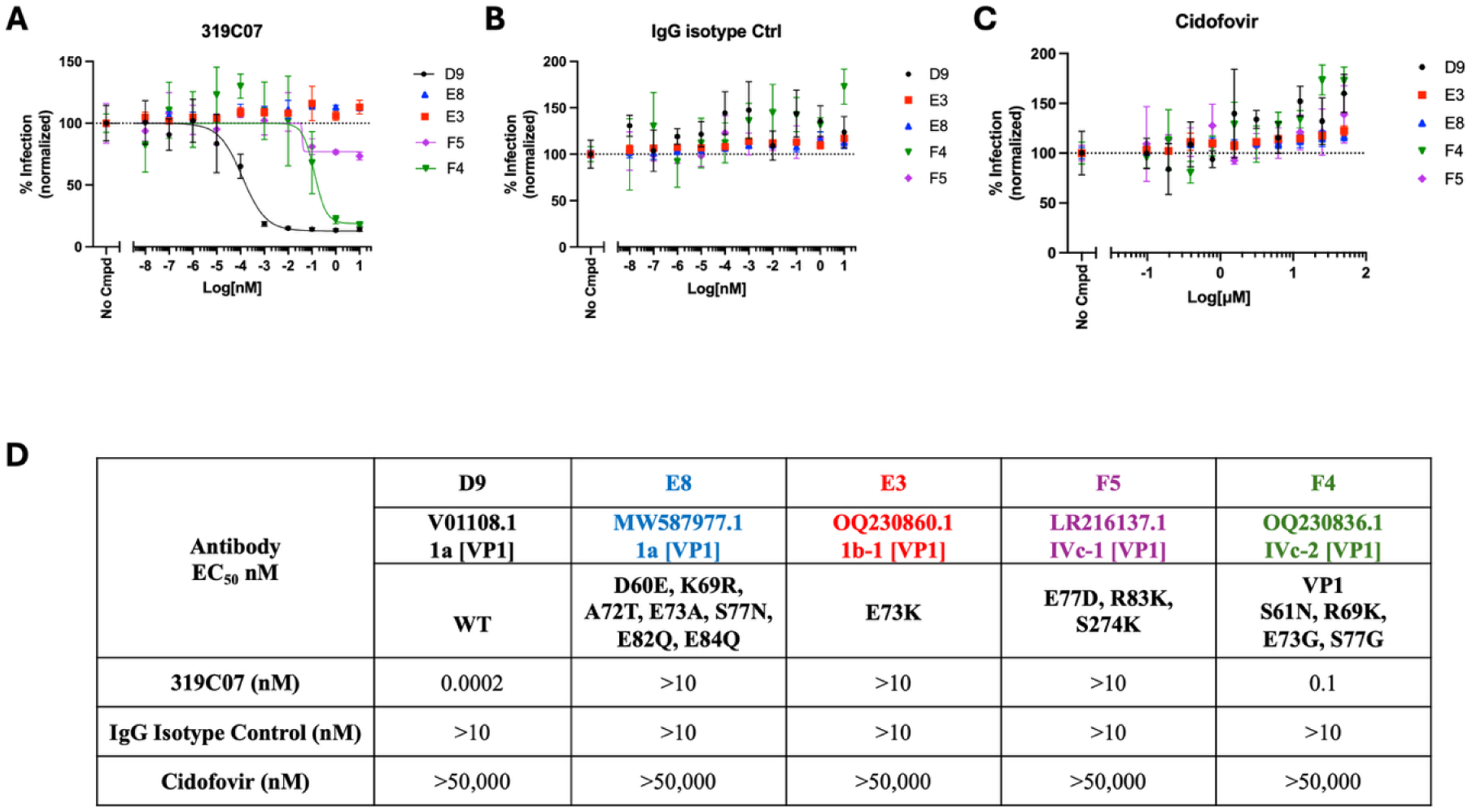
Neutralization potencies of monoclonal antibodies against BKPyV VP1 variants derived from clinical sequences. Dose response curves of mAb **A)** 319C07, **B)** an IgG isotype control and **C)** cidofovir, a nucleoside analog inhibitor of viral replication, respectively. Panel D shows the EC_50_ values representing neutralization potencies.

### **I.** Small molecule series with high affinity binding to both BKPyV and JCPyV VP1 support an antibody-independent path to targeting polyomaviruses

Given the immunologic plasticity of the BC-loop epitope, we explored alternative therapeutic modalities with the potential to access buried or sterically constrained regions. To assess whether conserved, non-immunogenic surfaces on VP1 could support small-molecule engagement, we evaluated a set of compounds from a custom-designed library focused on capsid-targeting ligands. One series representative, designated ORT-3198, exhibited high-affinity binding to recombinant BKPyV VP1 pentamers, with a dissociation constant (KD) of 1.3 nM measured by grating-coupled interferometry (Wave) and preserved high affinity binding to JCPyV VP1 (KD = 40 nM) (**Figure 9A and 9B**). Isothermal titration calorimetry (ITC) confirmed these interactions in an orthogonal readout of target-engagement. In a cell-based assay of BKPyV replication, ORT-3198 demonstrated antiviral activity against BKPyV with an EC_50_ of 4 µM and selectivity index of ∼10x, providing proof of concept of the first polyomavirus VP1 targeting small molecule (∼500 Da) (**Figure 9**). To our knowledge, ORT-3198 is the first drug-like small molecule shown to directly bind BKPyV VP1 with high-affinity and inhibit replication in primary human renal epithelium cells, complementing antibody-based anti-BKPyV strategies. While structural and mechanistic analyses of the binding site and optimization to close the affinity-activity window are ongoing, these findings establish that conserved VP1 surfaces may support both high-affinity small-molecule binding and antiviral function.

**Figure 9:**
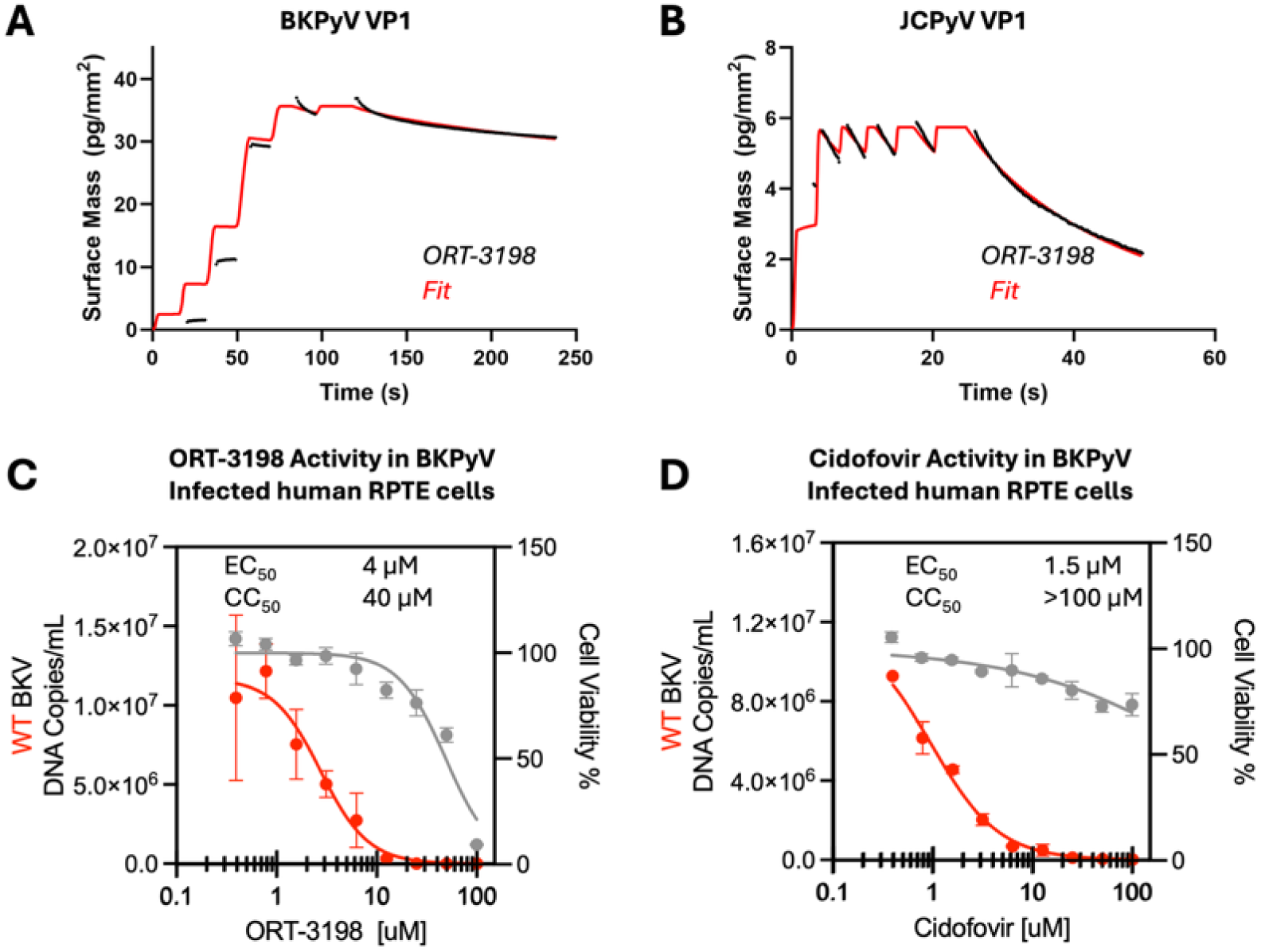
Discovery of a small molecule series targeting *BKPyV and JCPyV* VP1. Real-time binding of ORT-3198 to recombinant VP1 pentamers measured by grating-coupled interferometry (WAVE). Red traces are sensorgrams, black lines indicate model fits **(A-B)**, both experiments use multi-injection association formats but with different injection protocols. BKPyV VP1 data **(A)** was acquired with prolonged association pulses; JCPyV VP1 **(B)** with short, sequential association pulses; assay protocols differ, response amplitudes and kinetics not comparable. **C)** and **D)** BKPyV (genotype 1a) antiviral activity and cell viability dose response curves of ORT-3198 and cidofovir, respectively, in human RPTE cells.

## Discussion

BK polyomavirus (BKPyV) remains a major cause of allograft loss following kidney and bone-marrow transplantation, yet no approved antiviral therapies exist. Recent efforts have advanced two VP1-targeting monoclonal antibodies, MAU868 [15] and 319C07 (Potravitug) [16] into clinical evaluation, both demonstrating potent *in vitro* neutralization. However, whether this *in vitro* potency advantage translates into clinical efficacy is under investigation. Our comparative structural analysis of BKPyV VP1-antibody complexes led to the classification of VP1-targeting antibodies into two mechanistic classes, apical binders engage the receptor-binding cleft and make extensive contacts across the BC-loop, whereas lateral binders, such as MAU868, approach the pentamer periphery, contacting the EF-loop and avoiding the receptor cleft, resulting in limited BC-loop overlap (**Figure 4**). Our X-ray crystallographic analyses of apical binders, including 319C07 and functionally analogous nanobodies (VHH16/17) reveal that they make extensive contacts around the BC-loop and directly engage the sialic acid binding site mimicking glycan interactions with high surface complementarity (**Figure 3**). Although 319C07 and MAU868 exhibit comparable binding affinity to VP1 (data not shown), the superior *in vitro* neutralization potency of 319C07 likely reflects its ability to sterically block and functionally compete with receptor engagement through topological mimicry of the native glycan ligand.

To assess the stability of apical epitopes across circulating BKPyV variants, we analyzed 189 non-redundant, patient-derived BKPyV VP1 sequences from public datasets. This population-level analysis revealed substantial amino acid variation across exposed capsid loops, with the BC-loop exhibiting the highest density of nonsynonymous mutations, followed by the EF-loop (**Figure 5**). Focusing on the BC-loop, which forms the primary contact site for apical binders such as 319C07, we identified numerous substitutions across circulating variants within the antibody footprint (**Figure 6**). Functional testing of selected clinical variants revealed that several of these substitutions decrease or fully abrogate 319C07 binding (**Figure 7**). Notably, multiple escape-associated mutations correspond to residues previously implicated in persistent viremia in transplant recipients, designated non-controllers of BKPyV replication [22, 25, 26]. These findings suggest that even before therapeutic intervention, a meaningful fraction of patients may harbor variants with preexisting epitope incompatibility and confer resistance to receptor-cleft-targeting antibodies. More comprehensive VP1 sequencing of BKPyV VP1 in transplant patients will be essential to define the breadth of pre-existing epitope escape and to inform rational antibody design and trial stratification.

Longitudinal sequencing studies in transplant recipients have shown that VP1 mutations across the BC-loop and adjacent surface loops accumulate progressively in patients with persistent viremia [22, 25, 26]. These clinical observations are consistent with layered immune escape under sustained neutralizing antibody pressure. *In vitro*, we observed similar trajectories under sub-curative antibody concentrations: resistance selection with 336F07 and 319C07 led to the emergence of a D60 deletion and an E138Q substitution, respectively, while selection with 41F17 and MAU868 produced the G176V and T179I variants in BKPyV VP1. Although the full functional consequences of these surface-exposed mutations remain under investigation, their recurrence across experimental and clinical contexts suggests they represent potential incipient escape pathways. Some may serve as permissive intermediates, or stepping-stone mutations that lower structural or energetic barriers to future substitutions or compensate for associated fitness costs. Notably, the E138 substitution was observed in six patients in our analyses of population-level dataset and has been independently linked to APOBEC3-mediated editing [27, 34] underscoring its relevance across immune and intrinsic mutagenic pressures. This dynamic is further amplified by BKPyV’s unusually high substitution rate, among the highest reported for any DNA virus [30] and by the profound CD4⁺ T cell depletion induced by transplant immunosuppression, which dismantles a critical layer of immune containment and accelerates the emergence of antigenic variants. These findings indicate that once sub-curative pressure is applied in the setting of uncontrolled viremia, BKPyV can traverse accessible resistance pathways that erode antibody recognition over time. This underscores the importance of early intervention, before such escape trajectories are initiated, and highlights the additional limitations of reactive therapy with surface-loop targeting antibodies in the face of ongoing replication.

The susceptibility to escape-driven viremia is not uniform across transplant recipients. In most cases, BKPyV reactivation originates from the donor kidney, which often harbors a VP1 subtype distinct from the recipient’s pre-immune repertoire [35, 36]. This immunotype mismatch may delay the development of *de novo* neutralizing responses, creating a temporal gap during which the virus can replicate, disseminate, and evolve under weak or absent humoral control. Patients who remain viruric may benefit from partial cross-reactivity, while those who progress to viremia likely experience a failure of serologic adaptation. In this context, the virus encounters a permissive environment, one where antibody pressure is insufficient to clear infection, but sufficient to select escape. This is further supported by our population-wide sequence analysis, which revealed widespread variation across surface-exposed loops and highlighted not only the preexistence of functionally disruptive substitutions at multiple apical contact residues, but also the virus’s repeated use of these positions in resistance evolution, suggesting that structurally accessible escape routes are already embedded within circulating strain diversity. These observations support a model in which donor-recipient VP1 compatibility and the timing of neutralizing antibody emergence are key determinants of clinical trajectory and argue for integration of VP1 subtyping and baseline serology into trial and treatment stratification strategies.

Even in patients who mount timely or potent neutralizing responses, anatomical barriers may render those antibodies ineffective at the site of viral replication [37]. BKPyV replicates in renal tubular epithelial cells, an immunologically privileged compartment shielded from circulating IgGs [38]. Under physiological conditions, antibodies do not cross into the tubular lumen, and while transient permeability may occur during episodes of inflammation, such access is neither durable nor reliably present across patients. The neonatal Fc receptor (FcRn), while active in proximal tubule recycling and salvage of IgG [39], does not facilitate antibody trafficking into the interstitial space or lumen in a way that would enable sustained antiviral activity. As a result, antibodies, whether endogenous or exogenous, may fail to achieve virologic source control despite systemic presence. Their activity is confined to reducing viremia but not preventing viral entry or dissemination into naïve renal epithelial cells or eliminating replication in the intrarenal reservoir. This compartmentalized pattern of antibody pressure is mirrored in the clinical mutational landscape of nonsynonymous VP1 mutations, particularly in the BC-loop, which are consistently observed in viremic plasma but are absent from matched urine-derived isolates in patients with isolated viruria [22, 40]. These findings reinforce that immune selection operates in the bloodstream, where IgG concentrations are sufficient to exert pressure, but not within the tubular compartment where BKPyV replication persists. The virus thus replicates behind an immune firewall, sheltered from both humoral containment and antibody-based therapies. Once established in this niche, replication may continue unchecked despite the presence of circulating neutralizing antibodies, challenging the assumption that systemic antibody titers alone predict therapeutic efficacy.

Given the anatomical exclusion of antibodies from the renal parenchyma, the presumed mechanism of action for sialic acid-receptor-binding, BC-loop-targeting apical antibodies such as 319C07 warrants reexamination. In cell-based assays, neutralization is typically defined by an antibody’s ability to block virus-cell engagement, often via steric interference at the receptor-binding site or other entry-related mechanisms. For receptor-cleft-targeting antibodies like 319C07, this activity maps directly to structural mimicry of the sialylated glycan receptor and likely yields potent inhibition *in vitro*. However, this mechanism presumes direct access to the site of infection. *In vivo*, where antibodies are anatomically restricted from the site of replication, receptor competition and entry blockade cannot occur. Under these conditions, VP1-targeting antibodies may instead act peripherally, binding circulating virions shed from the kidney, promoting systemic clearance, or preventing reseeding of naïve epithelial targets, although BKPyV is known to replicate exclusively in the kidney. These downstream containment effects stand in contrast to neutralizing antibody therapies developed for SARS-CoV-2, RSV, or HIV, where viral replication occurs in compartments accessible to systemic IgG exposure. The observed reductions in plasma viral loads following VP1 antibody treatment, when present, likely reflect indirect containment of viremia rather than direct inhibition of viral replication.

Further mechanistic analysis is needed to clarify how VP1-targeting antibodies function *in vivo*, whether through hepatic clearance of immune complexes, modulation of FcRn trafficking pathways, or other systemic effects that diverge from classical receptor blockade. Without such understanding, *in vitro* neutralization potency remains an incomplete surrogate for therapeutic efficacy in the context of BKPyV VP1.

During the preparation of this manuscript, Phase II trial results for 319C07 (potravitug) in kidney transplant recipients with sustained BK viremia (median baseline ∼4.1-4.3 log₁₀ IU/mL) were publicly disclosed [41]. The trial reported no statistically significant difference in viral clearance (17.5% vs. 12.9%), a modest increase in the proportion of patients with ≥1-log viral load reduction (61% vs. 40.5%), and a non-randomized improvement in BKPyVAN histology despite higher baseline disease severity in the treatment group. These results are consistent with the constraints outlined above. Many circulating variants associated with persistent viremia harbored resistance mutations within the 319C07 binding epitope. In parallel, limited intrarenal antibody access likely prevented suppression of replication at the source. Sub-curative antibody exposure over 20 weeks could further enable variant evolution. Finally, many patients entered the trial with high viral loads, a context in which peripheral antibody activity is unlikely to meaningfully alter clinical trajectory. In a separate Phase II study, MAU868 was evaluated in kidney transplant recipients with low to moderate BK viremia, demonstrating higher rates of viral clearance (up to 75% <LLOQ) and ≥1-log reductions (60-70%), though without histologic endpoints or evidence of sustained clinical benefit [15]. In this setting with lower viral burden, peripheral containment mechanisms may appear more visible. However, neither MAU868 nor 319C07 achieved source control, reinforcing the broader limitation of VP1-targeting antibodies in the context of polyomavirus infections, which replicate within immune-privileged compartments. Together, these findings may explain the mismatch between high *in vitro* potency of VP1-targeting antibodies and their limited clinical efficacy.

To bypass the limitations of IgG-based therapies, we evaluated potent nanobodies which could possess enhanced renal penetration. However, like 319C07, the nanobodies tested, targeted the same receptor-cleft region and remained constrained by epitope-specific escape. This frames a mechanistic trade-off in which very high neutralization potency could be linked to engagement of highly mutable, receptor-cleft surfaces. This raises the possibility that nanobodies targeting conserved, non-cleft surfaces may have been deprioritized during selection due to lower apparent neutralization readouts, despite offering greater resilience to escape. Nanobodies can be readily reformatted into bi-paratopic or multivalent constructs, enabling combinatorial engagement of both potency driving and conserved surfaces. Such formats may mitigate escape risk by distributing binding energy across distinct topologies and broaden the therapeutic potential of nanobodies beyond single-epitope dependence.

In contrast, small molecules offer fundamentally distinct advantages: superior tissue penetration, the ability to bind buried or non-immunogenic capsid surfaces less prone to immune-driven variation, and the capacity to terminate viral replication through intracellular engagement. Our identification of high-affinity BKPyV VP1-binding compounds that retain high affinity binding with JCPyV VP1 underscores the feasibility of capsid-directed small-molecule antivirals (Figure 9). A series representative, ORT-3198 showed single digit micromolar EC_50_ value in an BKPyV infected human RPTE cell-based assay that was comparable to cidofovir. Additional optimization for enhanced antiviral activity, as well as the understanding of mode and site of binding could reveal opportunities for a new class of anti-polyomavirus agents. This approach may enable both therapeutic and prophylactic interventions across a broader range of viral titers and stages of infection, independent of host immune status or antibody access.

Several limitations of this study should be acknowledged. While we demonstrate that clinically observed VP1 variants disrupt antibody binding, additional neutralization profiling of these escape mutations is ongoing to fully characterize the affinity-activity relationship. The compartmental constraints of antibody access are inferred from pharmacologic and histologic literature but were not directly measured in this study. The relative contribution of cytotoxic T lymphocytes versus antibodies in BKPyV control has not been systematically quantified in transplant cohorts and was not assessed here; this distinction is critical to interpret whether persistent viremia reflects failure of antibody function, or of cellular immunity, or both. Lastly, our small-molecule findings represent a proof of concept and continued interrogation of their mode of action, and optimization will be required to translate high-affinity VP1 engagement into robust antiviral activity. Addressing these limitations in future studies will be critical to refining therapeutic strategies against BK and other polyomaviruses.

## Materials and methods

### X-ray crystallography of VHH16, VHH17 and 319C07

*Alpaca immunization and nanobody library generation:*VHH16 and VHH17 were generated under a contract with Eurogentec (Belgium). Two llamas were immunized with BK VP1.

PBMCs were isolated from blood on day 43 and subject to RNA isolation, quality assessment (18S and 28S rRNA) and reverse transcription into cDNA. IG-H cDNA fragments were amplified using specific primers annealing at the IGH leader sequence region and the CH2 region. The 700bp fragment representing VHH was excised and purified from gel and used as a template for nested PCR to introduce 5’ flanking *Sfi*I restriction site. The purified product was digested with *Sfi*I and *Eco91*I, ligated in frame with geneIII into pQ81 phagemid vector and transformed into *E. coli* TG1 by electroporation to generate the nanobody library of size ∼1x10^9^ and 100% insert frequency.

Following two rounds of panning using phages produced from the libraries, a concentration dependent enrichment was observed for BK VP1. Single clones from second round outputs were picked to create master plates. Periplasmic extracts were produced and screened for strong binding to BK VP1. Master plates were sequenced using Sanger sequencing and VHH sequences were clustered based on 90% CDRH3 homology. Multiple large clusters and unique sequences were identified. VHH16 and VHH17 were identified for further characterization.

*Vectors and cloning:* Gene fragments encoding the different serotypes of VP1 and a lysine substituted anti-ALFA-tag VHH (ALFAbodyK3) were cloned into an internal SUMO-tag fusion vector (pETSUK3), which includes an N-terminal His6 tag for Ni (II) chelate affinity purification. For the VP1 constructs for BLI an additional N-terminal ALFA-tag was also included in the construct after the SUMO sequence.

Both VHH16 and VHH17 was cloned into the *Nco*I-*Sma*I cloning site of the vectors from the pETCH family. These vectors included a PelB secretion sequence in the amino-terminal, and an uncleavable carboxyl-terminal His6 tag for Ni(II) chelate affinity purification.

*Recombinant protein production:*Antibodies were produced by GenScript turboCHOTM v2. Unless noted otherwise, the VP1 pentamers, VHHs and scFvs were produced as outlined below.

*Expression and purification of pentameric BK VP1 and ALFAbody:*All VP1 constructs were co-expressed with pRARE2LysS (Novagen), while the ALFAbody was co-expressed with a modified variant of pCyDisCo [42] in *Escherichia coli* T7 express cells (New England BioLabs, Ipswich, USA). Cells were induced at an optical density at 600 nm (OD600) of 2.0 with 100 µM and 500 µM IPTG respectively and incubated at 18 °C for 16 h. Cells were collected by centrifugation, resuspended in Resuspension Buffer (RB, 75 mM NaCl, 12.5 mM Imidazole, 50 mM Sodium Phosphate pH 7.5). The cell pellets were thawed and diluted with RB, supplemented with MgCl_2_ (2 mM) and DENERASE® (5,000 kU, >250 U/uL, c-LEcta GmbH, Leipzig, Germany) (0.5 uL/10 mL RB), and lysed by sonication (Sonifier SFX 150, Branson Ultrasonics, Bruckfield, USA).

For ALFAbody, sonication was repeated a total of 3 times (6 minutes, 5-second pulse on, 2- second pulse off at 55% amplitude). Lysate was cleared by a first centrifugation at 12,500 g for 30 minutes, followed by a high-speed centrifugation at 19,000 g for 30 minutes. The soluble fraction was loaded onto a 10 mL nickel-nitriloacetic acid (Ni-NTA) column (PureCube Ni-NTA Agarose, Cube Biotech, Monheim am Rhein, Germany) and subsequently washed with approximately 20 column volumes (CV) of wash buffer (WB, 250 mM NaCl, 12.5 mM Imidazole, 50 mM Sodium Phosphate, pH 7.5) by gravity flow. Proteins were then eluted in 3 mL fractions with elution buffer (EB, 250 mM, NaCl, 300 mM Imidazole, 50 mM Sodium Phosphate, pH 7.5). 1 µg of SUMO hydrolase per 100 µg of ALFAbody was added and then dialyzed against dialysis buffer (10 mM NaCl, 20 mM TRIS-HCl, pH 7) overnight at room temperature using a SnakeSkin Dialysis Tubing 3.5 K MWCO (Thermo Fisher Scientific, Waltham, USA). The dialysis buffer was refreshed the next morning, and dialysis proceeded for at least 1 hour. Cleaved ALFAbody was further purified by subtractive IMAC by using 10 mM NaCl, 12.5 mM Imidazole, 20 mM TRIS-HCl, pH 7 buffer, followed by cationic exchange chromatography (CEX) using a HiTrap SP HP column (Cytivia) using 0-40% gradient (Buffer A: 10 mM NaCl, 20 mM TRIS-HCl, pH 7, Buffer B: 1 M NaCl 20 mM TRIS-HCl pH 7) in 20 CV. The ALFAbody had a last clean-up step by size exclusion chromatography (SEC) using a Superdex S-75 pg column (Cytivia) using PBS buffer. For BLI, the ALFAbody was biotinylated using the manufacturer protocol of EZ-Link NHS-Biotin (Thermo Scientific, Waltham, USA)

For VP1 purification, sonication was repeated a total of 3 times (1 minute, 1-second pulse on, 1- second pulse off at 55% amplitude). Lysate was cleared by a high-speed centrifugation at 19,000 g for 30 minutes. The soluble fraction was loaded onto a 2 mL Ni-NTA column and subsequently washed with approximately 20 CV of WB by gravity flow. To speed up the purification process, on-column cleavage was performed, by adding 2 mL of WB to a capped column and 10 µg of SUMO hydrolase. The pentameric VP1 was left incubating with SUMO- Hydrolase for 1 hour at room temperature and 110 rpm. The cleaved VP1 was collected in the flow through by removing the cap of the column and adding extra 2 mL of WB to recover any cleaved VP1 from the void volume. The VP1 protein was then ammonium sulphate precipitated at 70% saturation, before a last purification step by SEC using a Superdex 200 increase 10/300 GL column (Cytivia) in PBS. The highest concentration fraction was used for BLI studies.

*Periplasmic expression and purification of VHHs:*VHHs (VHH16 and VHH17) were expressed in *Escherichia coli* T7 express cells (New England BioLabs, Ipswich, USA). Cells were induced at optical density of 2 at 600 nm (OD600) with 100 µM IPTG at 18 °C for 16 h. Cells were collected by centrifugation, resuspended in 1 mL per gram of pellet in TES buffer (200 mM TRIS-HCl, 0.5 mM EDTA, 500 mM Sucrose pH 8) and stored at -20 °C.

For lysis the cell pellets were thawed and incubated at 10 °C and 275 rpm for an hour. Then 2 mL per gram of pellet of 5 mM MgSO_4_ and 1 µL of Denerase (DENERASE® 5,000 kU, >250 U/µL, c-LEcta GmbH, Leipzig, Germany) were added. The cell pellet was then incubated at 10°C and 275 rpm for 1 to 2 hours. Lysate was cleared by a centrifugation at 12,500xg for 30 minutes. The soluble fraction was loaded onto a 2 mL nickel-nitriloacetic acid (Ni-NTA) column (PureCube Ni-NTA Agarose, Cube Biotech, Monheim am Rhein, Germany) and subsequently washed with approximately 20 CV of wash buffer (250 mM NaCl, 12.5 mM Imidazole, 50 mM Sodium Phosphate, pH 7.5) by gravity flow. Proteins were then eluted in 0.5 mL fractions with elution buffer (250 mM NaCl, 300 mM Imidazole, 50 mM Sodium Phosphate, pH 7.5). The eluted protein was ammonium sulfate precipitated to 70% saturation. The precipitated protein was spun down at 19,000xg for 30 minutes and resuspended in 0.2X PBS and desalted using a HiTrap desalting column (Cytivia) using PBS buffer. For Crystallization, VHH16 was further purified by size exclusion chromatography (SEC), using a Superdex S-200 column (Cytivia) in 150 mM NaCl, 10 mM TRIS-HCl, pH 7.5 buffer.

*Protein X-Ray crystallography:*VP1/ VHH16: An excess of VHH16 was mixed with BKPyV VP1 (residues 26-297) in a 1:2 molar ratio. and the complex was purified by SEC using a Superdex S-200 column (Cytivia) in 150 mM NaCl 10 mM TRIS-HCl pH 7.5. Crystals of VHH16/BKPyV VP1 were obtained at 8.3 mg/ml in the presence of 200 mM NDSB256 and using crystallization solution (30% (v/v) PEG 400 100 mM Sodium acetate/ Acetic acid pH 4.5 200 mM Calcium acetate) in a 1:1 ratio.

VP1/VHH17: An excess of VHH17 was mixed with BKPyV VP1 (residues 26-297) in a 1:2 molar ratio. and the complex was purified by SEC using a Superdex S-200 column (Cytivia) in 150 mM NaCl 10 mM TRIS-HCl pH 7.5. Crystals of VHH17/BKPyV VP1 were obtained at 10.2 mg/ml using crystallization solution Morpheus H7 (10% w/v PEG4000, 20% v/v glycerol, 0.1M MOPS/HEPES-Na pH 7.5, 0.02 M sodium L-glutamate, 0.02M DL-alanine, 0.02M glycine, 0.02M DL-lysine HCl, 0.02M DL-serine).

VP1/scFv: An excess of scFv-319C07 was mixed with BKPyV VP1 (residues 26-297) in a 1:1.25 molar ratio and the complex was purified by SEC using a Superdex S-200 column (Cytivia) in 150 mM NaCl 10 mM TRIS-HCl pH 7. Crystals of scFv-319C07/BKPyV VP1 were obtained at 6.4 mg/ml using crystallization solution (10% (w/v) PEG 6,000 100 mM Bicine pH 9) in a 1:1 ratio.

*Structure determination:* Crystals of VP1/VHH16, VP1/VHH17, VP1/scFv were cryoprotected with 20% v/v ethylene glycol and flash cooled in liquid nitrogen. Diffraction data was collected at Soleil synchrotron PIXII, for VP1/VHH16 and VP1/VHH17, and ALBA for VP1/scFV. All diffraction data was collected at 100K. Data was integrated using X (via XIA2), and integrated intensities were merged and scaled using programs from the CCP4i2 package [43]. The structures were solved by molecular replacement (MR) using models of VP1 (unpublished), VHHs (alfabuilder) and scFv, and PHASER [44]. We performed iterative cycles of model building and refinement using REFMAC [45] and manual model building was performed in Coot [46]. The quality of the final models was assessed by MolProbity and the CCP4i2 validation task.

### Bioinformatic analyses of BKPyV Sequences

Protein sequences for BK Polyomavirus 1 (BKVP1) were retrieved from the National Center for Biotechnology Information (NCBI) protein database. An initial query yielded a total of 3,048 BKVP1 protein sequences, encompassing both full and partial sequence annotations. The retrieved dataset underwent a rigorous curation process to ensure data quality and relevance.

First, sequences identified as artifacts or misannotated were removed, resulting in a refined dataset of 2,944 full and partial BKVP1 sequences. From this curated set, a subset of 923 BKV VP1 sequences was extracted that correspond to unique clinical isolates (based on genome accession IDs)[47, 48]. The unique protein sequences were aligned using MAFFT [49, 50].

Following alignment, the sequences were manually inspected using Jalview [51], with truncated sequences identified and removed. Finally, redundant sequences were eliminated using Jalview’s redundancy removal functionality, resulting in a final dataset of 190 non-redundant (NR) sequences, crucial for reducing computational burden while maintaining representative sequence diversity. Shannon entropy scores were generated within multiple protein sequence alignment to assess the sequence variability across BK VP1 region. For any position in the alignment, Shannon entropy (H) was calculated using the formula below:

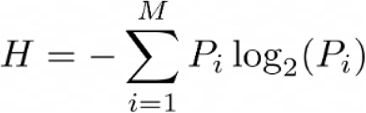

*where:*

*Pi: Probability (frequency) of amino acid type i at that position*.

*M: Total number of possible amino acid types (typically 20 for standard proteins). Logarithm is base 2, yielding entropy in "bits."*

The resulting scores were ranged from “lowest entropy (0 bits)” indicating complete conservation to “highest entropy [log_2_(20) or ≈4.32 bits]” indicating highest variability.

### Determination of binding affinity of nanobodies VHH16, VHH17 and mAbs 319C07 and 336F07 to BKPyV VP1 using biolayer interferometry (BLI)

*BLI assays and analyses:*For BLI we used a ALFAbodyK3 and the Biotin No weigh™ format EZ-link™ Sulfo-NHS-LC-LC-Biotin No weigh™ kit for immobilization. High-precision streptavidin biosensor tips were pre-incubated for 10 min in BLI buffer (20 mM Tris/HCl pH 8.0, 150 mM NaCl, 0.1% (w/v) bovine serum albumin, 0.02% (v/v) Tween-20). Tips were then dipped into BLI buffer for 60 s. Next, biotinylated, C-terminally tagged ALFAbodyK3 prepared in BLI buffer solution were immobilized to a thickness of ∼0.15 nm for 30 s. Next the tips were loaded with ALFA-tagged VP1 serotypes prepared in BLI buffer to a thickness of ∼0.5 nm for 60 s. Ligand-loaded tips were then dipped into wells containing BLI buffer for 60 s to determine a baseline. Tips were then transferred to solutions containing different antibodies or VHHs. Association was recorded for 300 s, followed by a 300 s dissociation step in wells containing BLI buffer. All binding sensorgrams were recorded on a forteBIO OctetRED96 instrument (Sartorius). A reference sensor was included and used to subtract background noise. Data were normalized to the baseline step and aligned to the dissociation point using the Octet data analysis software. Data was plotted using PRISM sofware (GraphPad).

### BKPyV pseudovirus reporter assay

*Preparation of BK Polyomavirus Pseudoviruses:* Pseudovirions were produced as previously described by Buck and colleagues [33] . 7x10^5^ HEK293TT cells were plated in a T75 cell culture flask one day prior to transfection. The following day, cells were transfected (6) with a mixture of Lipofectamine 2000 (Thermo Fisher Scientific) and four plasmids encoding for VP1 (pIaw, 9.5µg), VP2 (ph2b, 9.5µg), VP3 (ph3b, 9.5µg), and Luciferase (pNL1.3.CMV, Promega, 9.5µg). To create pseudovirions with various VP1 serotypes, the VP1 coding region of pIaw was exchanged with the desired serotype sequence. 5-hours post transfection, transfection media was replaced with cell culture media (DMEM containing 10% FBS, 1X NEAA, and 1X Glutamax; DMEM-10), and cells were kept for an additional 48 hours at 37°C. Cells were harvested and washed with DPBS (Thermo Fisher Scientific), cell lysis and maturation of the resulting pseudovirions occurred by sequential addition of 2 U/mL Neuraminidase V (Sigma Aldrich), 50mM Tris pH 8.0, 0.5% Brij-58 (Sigma Aldrich), 1:1000 dilution of RNase Cocktail Enzyme Mix (Thermo Fisher Scientific), and 25mM Ammonium Sulfate pH 9.0 then incubated at 37°C for 20-24 hours. After maturation, pseudovirions were separated from cell debris by multiple rounds of centrifugation and washing. The resulting cell pellet underwent two freeze-thaw cycles and centrifugation, after which all supernatants were pooled. Pseudovirions in the pooled supernatants were purified from other cellular proteins using gel filtration (2% BCL Agarose Bead Standard (50-150µm), Agarose Bead Technologies). Gel filtration fractions were tested for the presence of encapsidated DNA prior to pooling using the Quant-iT PicoGreen dsDNA Assay Kit (Thermo Fisher Scientific). PsV titers were determined by TCID_50_ using the Spearman- Karber algorithm. Briefly, HEK293TT cells were seeded at a density of 10,000 cells/well in a 96-well plate one day prior to infection. PsV stocks were serially diluted in the 96-well plate and incubated for 3 days at 37°C. Infectivity was measured via quantification of secreted Nano- luciferase using the NanoGlo Luciferase Assay (Promega) in the VICTOR Nivo plate reader (Revvity).

*Neutralization of BK Polyomavirus Pseudoviruses:*HEK293TT cells were seeded at a density of 10,000 cells/well in cell culture medium (DMEM-10) one day prior to assay and left to adhere/double overnight. The following day, 10 serial dilutions of either antibody or compound were prepared in cell culture medium. PsV inoculum was prepared for a final MOI of 0.05.

Antibody/compound dilutions and BKPy PsV were mixed and incubated for 1 hour at 37°C prior to addition to the cells. After 1 hour, all media was removed from the cells, and 100uL of the treatment/PsV mixture was added to each well. The plate was incubated for 3 days at 37°C. After 3 days, secreted Nano-luciferase was quantified by the NanoGlo Luciferase Assay (Promega).

Briefly, 40 µl of cell supernatant was transferred to a white, flat bottom 96-well plate and mixed with 40 µl of Nano-Glo® Luciferase Assay Buffer and Substrate mix. After a 5 to 10-minute incubation at room temperature, luminescence was quantified by a VICTOR Nivo plate reader (Revvity). Luminescence was analyzed with an integration time of 1s. BKPy PsV infection was first blank corrected (no infection) and then normalized to infection alone (set to 100). Data and IC_50_ values were determined through analysis in GraphPad Prism 10.

### BKPyV infection assay in human RPTE cells

Primary human renal proximal tubule epithelial (HRPTE) cells (ATCC) were cultured in Epithelial Cell Medium (EpiCM; Innoprot) supplemented according to the manufacturer’s instructions. Cells were maintained at 37°C in a humidified incubator with 5% CO₂ and used for experiments up to passage six. To evaluate antiviral activity of mAbs, nanobodies, or cidofovir, HRPTE cells were seeded at a density of 7,500 cells per well in 96-well plates 18 hours prior to infection. BKPyV (Genotype 1a Dunlop strain ATCC VR-837) was diluted in pre-warmed EpiCM to achieve multiplicity of infection (MOI) of 0.01. The virus was pre-incubated with serial dilutions of test mAbs/nanobodies/cidofovir, in triplicates for 1 hour at 37°C. At the start of the assay, the culture medium was removed, and 100 µL of the virus-compound mixture was added to each well. Plates were incubated at 37°C/5% CO₂ for 1 hour, after which the inoculum was removed and replaced with fresh EpiCM containing the corresponding compound dilutions. Cells were incubated for seven days at 37°C/5% CO_2_, after which viral replication was assessed by quantifying viral genome copy numbers in the supernatant via qPCR and a standard curve. EC_50_ of compounds/mAbs/nanobodies was determined as a 50% reduction in viral copy number from the untreated (DMSO treated) control using a 4-parameter curve fitting algorithm.

To evaluate compound cytotoxicity independently of viral infection, HRPTE cells were seeded as described above. The following day, cells were treated with serial dilutions of the test mAbs, nAbs, or small molecules in EpiCM medium without virus in triplicates. After a 7-day incubation at 37°C, cell viability was assessed using the XTT assay (GoldBio, Cat# X-200-1) according to the manufacturer’s instructions. CC_50_ of compounds/mAbs/nAbs was determined as a 50% reduction in cell viability from the untreated (DMSO treated) control using a 4-parameter curve fitting algorithm.

## Acknowledgements

The authors would like to thank Maria Martin for providing operational and administrative support in coordinating and managing external studies.

## Notes

### Competing Interest Statement

OA, CUN, AB, CCS, FHM, SDG, EED, SJR, MC, NM, SDW and AHM are employees of Pledge Therapeutics and/or equity holders in Orthogon Therapeutics. VCC serves as a clinical advisor to Orthogon Therapeutics.

